# PreciCE: Precision engineering of cell fates via data-driven multi-gene control of transcriptional networks

**DOI:** 10.1101/2024.11.04.621938

**Authors:** Jens P. Magnusson, Yusuf Roohani, Daniel Stauber, Yinglin Situ, Paloma Ruiz de Castroviejo Teba, Rickard Sandberg, Jure Leskovec, Lei S. Qi

## Abstract

The directed differentiation of stem cells into specific cell types is critical for regenerative medicine and cell-based applications. However, current methods for cell fate control are inefficient, imprecise, and rely on laborious trial-and-error. To address these limitations, we present a method for data-driven multi-gene modulation of transcriptional networks. We develop bidirectional CRISPR-based tools based on dCas12a, Cas13d, and dCas9 for simultaneously activating and repressing many genes. Due to the vast combinatorial complexity of multi-gene regulation, we introduce a machine learning-based computational algorithm that uses single-cell RNA sequencing data to predict multi-gene perturbation sets for converting a starting cell type into a desired target cell type. By combining these technologies, we establish a unified workflow for data-driven cell fate engineering and demonstrate its efficacy in controlling early stem cell differentiation while suppressing alternative lineages through logic-based cell fate operations. This approach represents a significant advancement in the use of synthetic biology to engineer cell identity.

## Introduction

Synthetic biology applies engineering principles to build biological systems. Recent advances in the field have been driven by the convergence of powerful computational tools, large biological datasets, DNA synthesis and sequencing technologies, and novel gene perturbation tools (e.g., CRISPR). Regenerative medicine and other cell-based applications could benefit tremendously from a synthetic biology-based approach where cell identity is rationally engineered in a data-driven way. However, until now, integration of the necessary computational and experimental tools has been lacking. Here, we sought to bring all these elements together into a single, unified method for cell fate control.

Cell identity can be controlled through small molecules, modulation of transcription factors, or CRISPR-activation (CRISPRa)^1–3^. Such strategies often rely on activating one or a few master-regulator genes, which can execute broad perturbations in transcriptional networks. But broad network perturbations can cause imprecision and heterogeneity on the single cell level – major challenges in stem cell engineering applications. Precision cell-fate control (defined as a high differentiation efficiency and low heterogeneity) may instead require careful sculpting of transcriptional networks. We hypothesized that simultaneous activation and repression of multiple genes can precisely fine-tune transcriptional networks, thus inducing desired cell states and blocking undesired ones.

Simultaneous gene activation and repression has been demonstrated with CRISPR-dCas9 technology in human cells^4–9^. With few exceptions^9^, however, these approaches have been limited to a “one-gene-up, one-gene-down” scheme. Downregulating one gene while upregulating another can indeed improve cell fate programming efficiency^6^ but has limited potential to reprogram the vast transcriptional networks that underlie cell fate commitment. New technology is needed that enables users to perform multi-gene activation and repression on a large scale.

Given such multi-gene control technology, however, how does one know which genes to modulate? Currently, empirical knowledge, intuition, and trial-and-error experimentation is required. Instead, we think precision cell-fate control must be data-driven. For example, large repositories of single-cell RNA sequencing (scRNA-seq) data exist and could serve as instruction manuals for cell fate conversion. Algorithms do exist that use such data to predict transcription factor perturbations^10–18^. Crucially, however, none of these predicts multi-gene sets for activation and repression, and they are therefore not suitable for precision control of multi-gene transcriptional networks.

To address these challenges, we developed a method for data-driven cell reprogramming via engineering of multi-gene transcriptional networks, which we call PreciCE (<u>Preci</u>sion <u>C</u>ell-Fate <u>E</u>ngineering, pronounced “precise”) (**Fig. 1**). PreciCE consists of two parts: computation and execution. The computation part consists of our new algorithm (“The PreciCE algorithm”), which translates scRNA-seq datasets of desired starting and target cell states into ‘genetic instructions’ for cell fate programming consisting of multi-gene perturbation sets. The execution part consists of three new CRISPR-based dual-Cas tools, each capable of performing simultaneous multi-gene activation and repression in human cells (“The PreciCE toolbox”). When combined, these technologies interface seamlessly to create a single workflow for data-driven precision cell-fate engineering. We show that (1) data-driven multi-gene perturbation enables higher precision than single-gene perturbation, (2) simultaneous up- and downregulation further enhances precision by actively blocking undesired cell states, (3) the unified integration of computational prediction and experimental tools brings synthetic biology to cell fate control, transforming cell differentiation into a precise engineering discipline.

**Figure 1.**
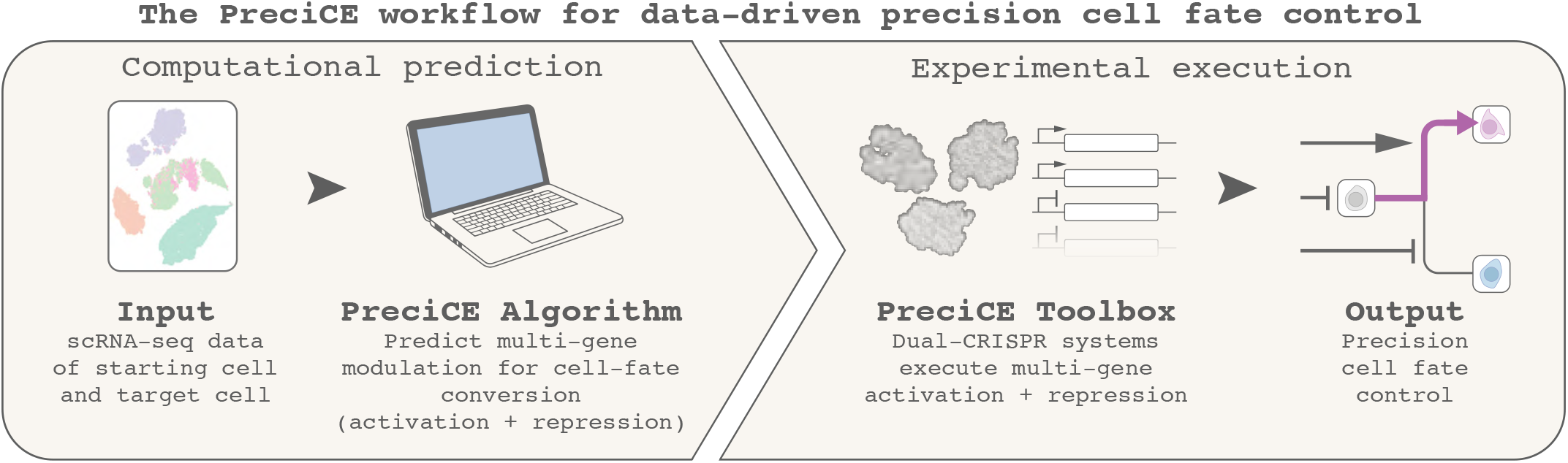
**PreciCE (<u>Preci</u>sion <u>C</u>ell-Fate <u>E</u>ngineering)**, a method for data-driven precision cell- fate programming. ScRNA-seq datasets (e.g., publicly available) feed into our PreciCE algorithm that converts these data into genetic instructions for cell fate programming via simultaneous up- and downregulation of multiple genes. To execute these complex genetic instructions, our PreciCE toolbox consists of three new dual-CRISPR systems for simultaneous multi-gene activation and repression. When combined, this workflow enables rational, data-driven specification of precise cell types while simultaneously repressing undesired, contaminating cell types for improved stem cell-based applications.

## Results

### The PreciCE toolbox: Three CRISPR-based systems for the simultaneous up- and downregulation of many genes

CRISPR arrays enable targeting multiple genes in a compact way^19^. We first validated that a recently reported *Lachnospiraceae bacterium* hyperactive dCas12a (hyperdCas12a), with its guide RNAs (gRNAs) encoded on a single CRISPR array, can mediate multi-gene activation (30 genes) (**Fig. S1A-B**), with high co-modulation of target genes (**Fig. S1C-D**) and only a moderate drop in CRISPRa efficiency with longer arrays (**Fig. S1E-F**). CRISPR arrays can be rapidly designed, synthesized, and assembled^20^. We therefore sought to expand the use of such arrays for simultaneous activation and repression, and developed three dual-Cas systems, all combining hyperdCas12a with another Cas protein for multiplexed up- and downregulation.

### A CRISPR-Cas13d/dCas12a hybrid array enables multi-gene activation and repression

We combined hyperdCas12a-miniVPR^21^ for programmable transcriptional activation with *Ruminococcus flavefaciens* Cas13d (Rfx-Cas13d)^22^ for targeted mRNA destruction. We chose this combination because both dCas12a and Cas13d can process their own CRISPR arrays^22,23^. Since the gRNA repeat sequences of Cas12a and Cas13d differ greatly (**Fig. S2A**) and are not recognized by each other’s Cas protein (**Fig. S2B-E**), we encoded their gRNAs on a single “hybrid” CRISPR array. We optimized multiple array architectures and parameters, focusing mostly on Cas13d optimization, as we and others have previously optimized CRISPR-Cas12a arrays^20,24^. We tested the effects of polymerase II (Pol. II) versus Pol. III promoters (**Fig. S2F-J**) in combination with Cas13d nuclear localization or export signals (**Fig. S2F-G**). We varied the position and amount of gRNAs on the array (**Fig. S2K-O**). And we confirmed that Rfx-Cas13d has collateral activity proportional to its target gene’s expression level^25^, though this effect could be removed by titering down expression of Cas13d’s doxycycline-inducible target gene GFP **(Figs. S2P-S, S3F**).

With our optimized hybrid array design (**Fig. 2A**), we first simultaneously downregulated one gene (GFP) and upregulated another gene (CD9), as measured by flow cytometry (**Fig. 2B**). Next, we demonstrated upregulation of three genes [*CD9↑ IFNG↑ IL1RN↑*] and simultaneous downregulation of three genes [*GFP↓ HRAS↓ SMARCA4↓*] in HEK293T cells, as measured by RT-qPCR (**Fig. S2T-U**) and scRNA-seq^26^ (**Figs. 1C, S3A-E),** with no detectable collateral effects (**Fig. S3F**). Thus, this design enabled multi-gene activation and repression using a single Pol. III-transcribed CRISPR array.

**Figure 2.**
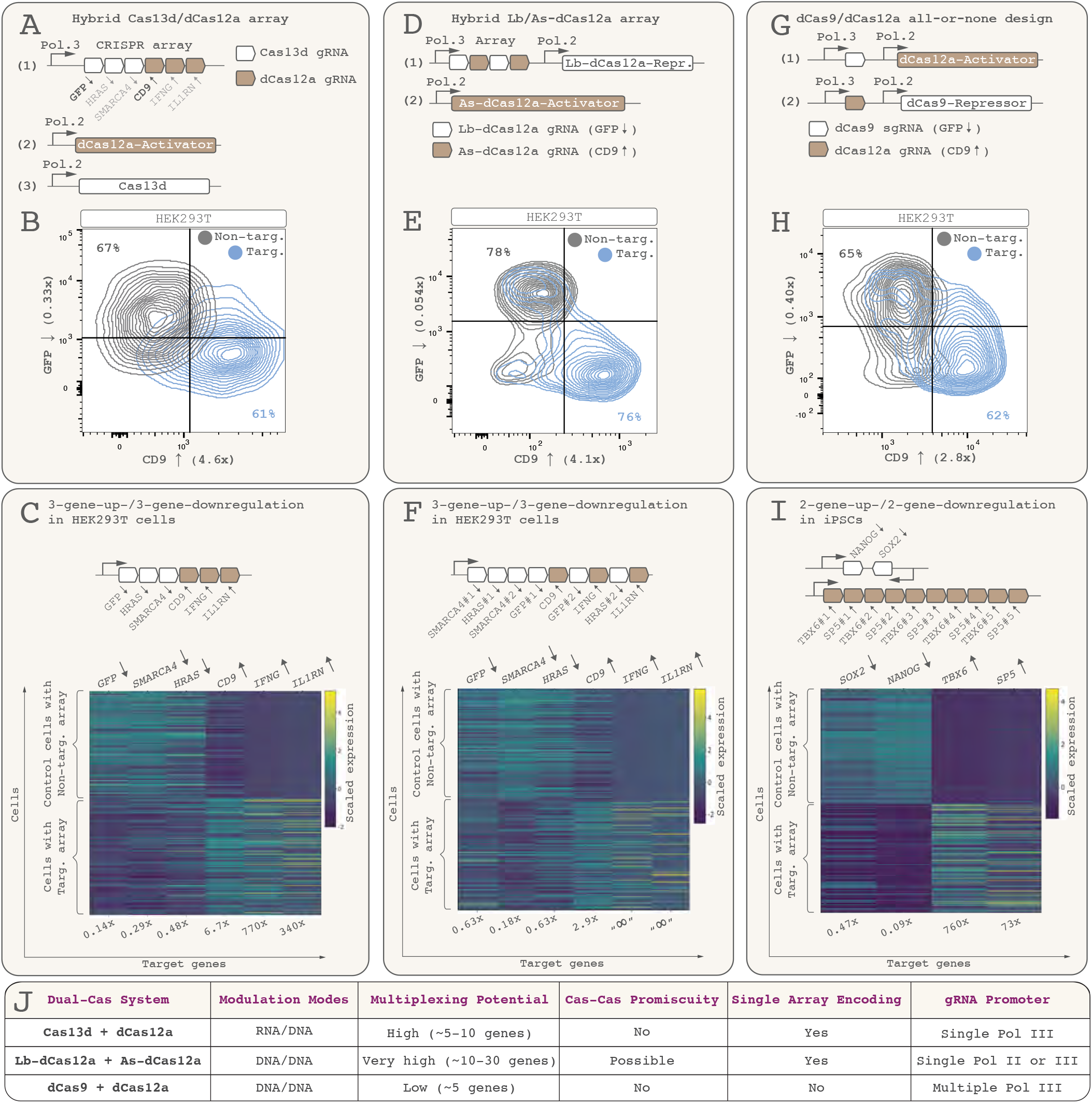
The PreciCE toolbox: Three CRISPR–based architectures for simultaneous activation and repression of multiple genes (**A**) A hybrid CRISPR-Cas13d/dCas12a array encodes Cas13d and dCas12a gRNAs on a single transcript. (**B**) When transfected with a dCas12a-miniVPR activator and Cas13d, this enables simultaneous upregulation of one gene (CD9) and RNA-targeted downregulation of another gene (constitutively expressed GFP) as measured using flow cytometry. (Numbers in axis legends represent expression ratio between targeting and non-targeting arrays, i.e., fold-change). (**C**) A scRNA-seq heatmap (Rows: cells, columns: genes) shows simultaneous downregulation of three genes [*GFP↓ HRAS↓ SMARCA4↓*] and upregulation of three genes [*CD9↑ IFNG↑ IL1RN↑*] in HEK293T cells (lower half of heatmap) compared to baseline (non-targeting gRNAs; upper half). Heatmap color scale spans the maximum and minimum expression value for each gene. Note that the fold-change metric is heavily affected by baseline expression level. (**D**) A hybrid CRISPR Lb-dCas12a/As-dCas12a array is encoded along with Lb-dCas12a-KRAB on a single construct. (**E**) It enables simultaneous gene repression (GFP) and activation (CD9) when co-transfected with As-dCas12a-miniVPR. (**F**) ScRNA-seq data showing simultaneous modulation of six genes, in this case by encoding two unique gRNAs for each Lb-dCas12a target gene. (**G**) A dCas9/dCas12a all-or-none design encodes the gRNAs for each Cas protein on the construct of the other Cas protein. Only cells that take up both constructs perform any gene modulation, preventing partial modulation and heterogeneity with plasmid delivery (e.g., transfection). (**H**) Upregulation of CD9 and downregulation of endogenous GFP, shown by flow cytometry. (**I**) ScRNA-seq data of iPSCs repressing two transcription factors [*SOX2↓ NANOG↓*] and activating two others [*TBX6↑ SP5↑*]. (**J**) Distinguishing features of the three multi-gene control systems. *Modulation modes*: whether gene modulation occurs on the RNA or DNA level. *Multiplexing potential*: The rough number of genes that can be targeted with the presented designs. *Cas-Cas Promiscuity*: Whether the two Cas proteins interfere with each other. *Single Array Encoding*: Whether gRNAs from both Cas proteins can be encoded on a single hybrid CRISPR array. *gRNA Promoter*: Recommended promoters for gRNAs based on our experimental data.

### A CRISPR Lb-dCas12a/As-dCas12a hybrid array enables activation and epigenetic repression of many genes

Some cell engineering applications may benefit from epigenetic repression, which can be more long-lasting than Cas13d-based RNA-level repression^27,28^. And because Cas12a’s gRNAs work when expressed under a Pol. II promoter, a dual-Cas12a hybrid system could enable very long CRISPR arrays and increased multiplexing capacity. So we sought to use two dCas12a variants for simultaneous activation and repression. We used engineered, nuclease-deactivated variants of *L. bacterium* Cas12a (*hyperdCas12a*^21^, subsequently denoted Lb-dCas12a) and *Acidaminococcus species* Cas12a (*enhanced As-dCas12a*^29^, denoted As-dCas12a).

We found that Lb-dCas12a and As-dCas12a can process and use each other’s gRNAs (**Fig. S4A-C**)^30^, with As-dCas12a being especially promiscuous (**Fig. S4B-C**). They also recognize the same protospacer-adjacent motif (PAM) sequence (TTTV^31^). Interestingly, though, when we used an engineered gRNA repeat sequence for As-dCas12a^32^ (**Fig. S4A-C**) and used the less promiscuous Lb-dCas12a for repression (**Fig. S4D-F**) and encoded the hybrid As/Lb-dCas12a gRNA array on the Lb-dCas12a construct (**Fig. 2D**), we achieved greatly improved orthogonality. Indeed, we simultaneously activated endogenous CD9 and repressed GFP in HEK293T cells (**Figs. 2E, S4G-H**), with no sign of promiscuous interference (**Figs. 2E, S4G**). We further achieved simultaneous repression of three genes [*GFP↓ HRAS↓ SMARCA4↓*] and activation of three genes [*CD9↑ IFNG↑ IL1RN↑*] using this system in HEK293T cells, as measured by scRNA-seq (**Figs. 2F, S4I-K**).

### A dCas9/dCas12a two-construct architecture ensures all-or-none up- and downregulation

In cell fate programming applications, a heterogeneous mix of cell states may arise if some cells fail to modulate all target genes. Therefore, we sought to devise a CRISPR architecture that would ensure all-or-none multi-gene regulation, especially for experimental settings prone to inefficient uptake and delivery. We devised a two-construct system with a dCas12a-miniVPR activator and a dCas9-KRAB repressor (**Fig. 2G**). We encoded the dCas12a gRNAs on the same plasmid as the dCas9 gene, and the dCas9 gRNAs on the same plasmid as the dCas12a gene. Thus, only cells co-expressing both constructs would simultaneously perform activation and repression (a logical AND gate), whereas cells that expressed only one of the constructs would experience no gene modulation. Indeed, co-transfection of these constructs (**Fig. S5A-C**) into GFP-expressing HEK293T cells executed simultaneous upregulation of one gene (CD9) and downregulation of another gene (GFP) (**Figs. 2H, S5D-E**). No gene was modulated in cells that only took up one of the constructs. We next simultaneously activated two transcription factors (*TBX6*, *SP5*) and repressed two others (*SOX2*, *NANOG*) in human induced pluripotent stem cells (iPSCs), as measured by scRNA-seq (**Figs. 2I, S5F-H**). Thus, this design allowed simultaneous activation and repression with minimal cell-to-cell heterogeneity.

Taken together, we developed three CRISPR-based systems for the simultaneous activation and repression of multiple genes. Each system has unique advantages and disadvantages (**Fig. 2J**), enabling users to choose the system that best matches their design specifications.

### Mechanisms of multi-gene regulation

We asked, using these multi-gene control systems, do individual cells modulate all target genes or does each cell only modulate some subset? Using a Wilcoxon signed rank test on scRNA-seq data (**Methods**), we found that many individual cells modulate all target genes (**Fig. S6A-C**). But the efficiency of multi-gene modulation was roughly equal to the multiplication product of how efficiently each individual target gene was modulated (**Fig. S6D-F**): A poorly performing gRNA reduces not only the modulation magnitude per cell but also the percentage of modulated cells^20^ (**Fig. S1E-F**). Thus, if some target genes are inefficiently modulated, multi-gene modulation may become patchy. For example, using our dCas9/dCas12a system (**Fig. 2G**), *NANOG* repression was more efficient than *SOX2* repression (**Figs. 2I, S5G**). So some cells repressed *NANOG* but not *SOX2* (**Fig. S6G-I**). Importantly, though, cells that modulated one gene were not less likely to simultaneously modulate another gene (**Fig. S6D-F**). In fact, cells were slightly more likely to modulate *all* target genes than subsets (**Fig. S6D-F**). We did find, though, that gene modulation efficiency was less efficient the more genes were targeted, consistent with dilution of available dCas protein (**Fig. S6J**). We later found that this effect could be counteracted by increasing the expression level of dCas genes (**Fig. S12A-B**).

Transcriptional bursting can be analyzed using Smart-seq data^33^, but how bursting is affected by CRISPR-based gene modulation is not known. Interestingly, for the target genes we studied, we found that CRISPR interference (CRISPRi, using dCas9-KRAB and dCas12a-KRAB) primarily acted by reducing transcriptional burst frequency, while burst size surprisingly increased slightly, possibly as a compensatory mechanism (**Fig. S7A-B**). CRISPRa (with dCas12a-miniVPR) primarily increased burst size (**Fig. S7C-D**). And we found evidence that dCas12a-miniVPR binding can shift the transcriptional start site slightly downstream, possibly due to steric effects (**Fig. S7E**). Cas13d-mediated repression was caused by a reduction in burst size but not frequency, consistent with Cas13d’s role as an mRNA-targeting enzyme (**Fig. S7F**). Interestingly, Cas13d-cleaved transcripts lingered in the cell long enough to be detectable in our scRNA-seq dataset, including in the form of partial transcripts missing the first few exons. (**Fig. S7G-I**). Finally, Pol. III-transcribed CRISPR arrays were poly-adenylated and captured in our sequencing libraries (**Fig. S7J-K)**, with possible implications for future multiplexed CRISPR screening.

### The PreciCE algorithm: A machine learning multi-gene prediction tool for data-driven cell fate conversion

To make precision cell fate control data-driven, we developed an algorithm that uses scRNA-seq data of a desired starting and target cell type as input (**Fig. 3A**). We call our model the PreciCE algorithm, and it operates as follows (**Methods**). First, it reconstructs transcriptional networks underlying these cell types (**Fig. 3B**. Users can choose to reconstruct transcriptional networks purely based on the input data or use a pre-generated network). It then trains a predictive model of gene expression following a combinatorial genetic perturbation by parameterizing the network edges. This model is then inverted to identify which transcription factors should be simultaneously activated and repressed to achieve an optimal transition from the starting cell state to the target cell state (**Fig. 3C**). To demonstrate, we used the PreciCE algorithm to predict a gene perturbation set for converting pluripotent stem cells (starting state) into mesoderm cells (target state), using a publicly available scRNA-seq data as input^34^. The algorithm’s output consists of a ranked list of transcription factors most likely to convert the starting state into the target state (**Fig. 3D**). This ranking is cumulative: each transcription factor should be perturbed together with all higher-ranking transcription factors. Each gene has a directionality of perturbation (upregulation: “↑”, downregulation: “↓”). A Precision Score (**Fig. 3D**) represents the algorithm’s estimate of how close the suggested perturbation will bring the starting state to the target state, and it increases cumulatively as more genes are included in the perturbation set. Each gene’s ranking reflects its overall effects on the transcriptional network – not merely its differential expression magnitude or how many downstream target genes it directly regulates (**Fig. 3E**). For example, *MIXL1* (excluded) is predicted to regulate more genes than *TBX6* (included). But *MIXL1* is not top-ranked because it is predicted to be modulated as a secondary effect when *TBX6* is modulated. As the Precision Score levels off, further perturbations are predicted to make little difference to the outcome.

**Figure 3.**
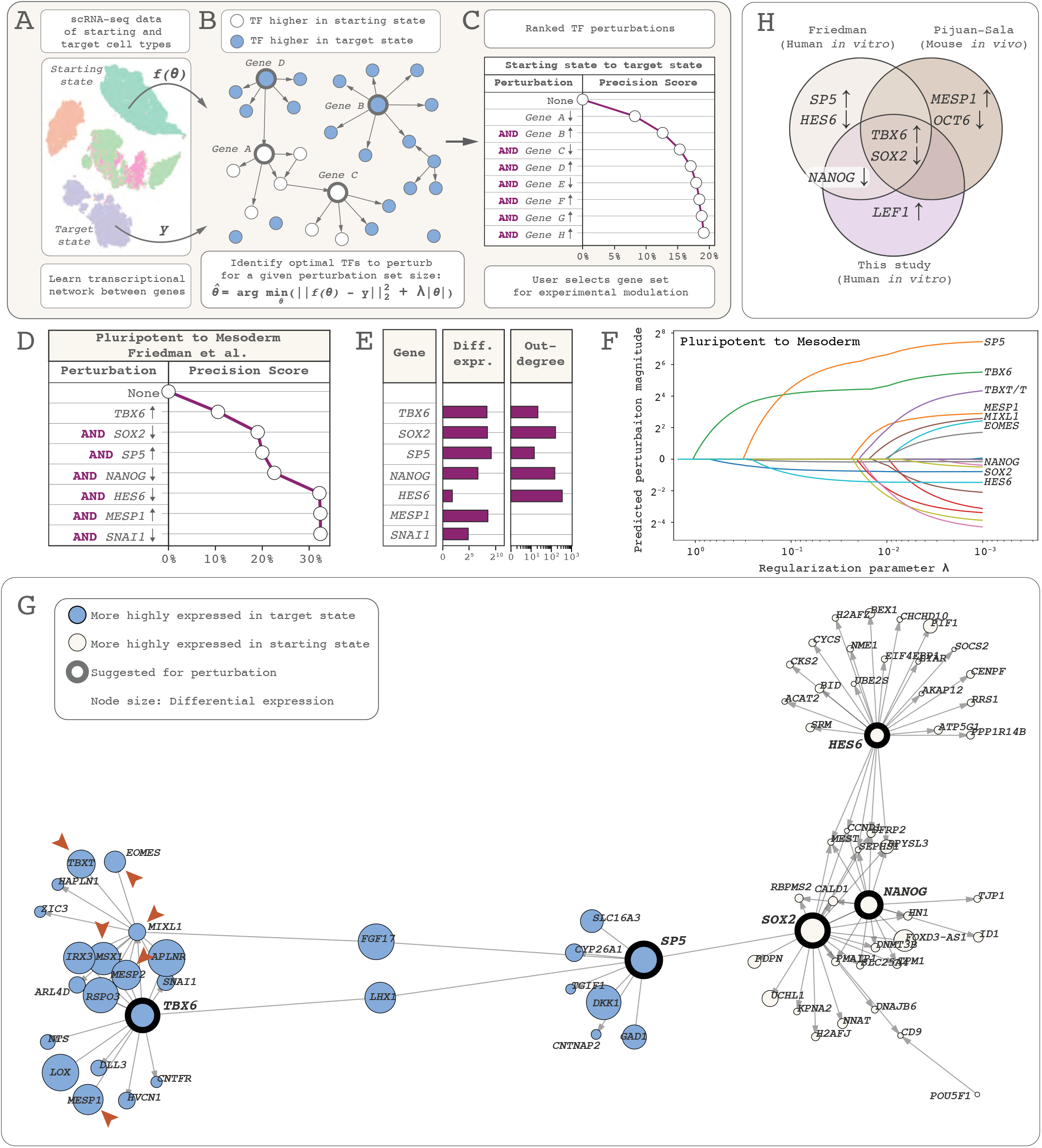
The PreciCE algorithm: A computational tool for data-driven cell fate conversion. (**A**) The PreciCE algorithm uses scRNA-seq data of a desired starting cell type and target cell type as input. (**B**) It reconstructs an underlying transcriptional network of differentially expressed transcription factors. Using this network, it identifies a set of transcription factors to modulate (up- and downregulation allowed), for most effectively destabilizing the starting cell state and activating the target cell state. (**C**) The output consists of a ranked list of transcription factors to activate (“↑”) or repress (“↓”). This list is accompanied by a Precision Score – an estimate of how similar the perturbed starting state will be to the target state. The list of transcription factors is cumulative: for each transcription factor down the list, the Precision Score reflects how efficient the cell state conversion will be when that transcription factor is perturbed together with all higher-ranked transcription factors. Based on where the Precision Score curve levels out, users can decide how many of the top-ranked transcription factors to modulate experimentally. In this example, a reasonable modulation gene set is [*Gene A↓ Gene B↑ Gene C↓ Gene D↑*]. (**D**) Predicted gene perturbation set for converting pluripotent stem cells into cardiogenic mesoderm. (**E**) The model ranks genes based on their overall network effects and not merely for being highly differentially expressed or regulating many genes (out-degree). (**F**) The PreciCE algorithm varies the constant λ iteratively and registers how the predicted perturbation effect changes (i.e., which transcription factors are chosen to be perturbed). This enables gene ranking when reading this graph from left to right. (**G**) The computationally reconstructed network of differentially expressed genes for this experiment, visualized using Observable, an online tool. Thick borders highlight genes suggested for perturbation. Node size represents differential expression magnitude between starting and target cells. Disconnected nodes (n=99) are excluded for clarity. Note that *TBX6* is predicted to directly or indirectly regulate many mesoderm-associated transcription factors (arrowheads; e.g., *MESP1*, *MESP2*, *MIXL1*, *EOMES*, *MSX1*, *TBXT*). (**H**) Across three different scRNA-seq datasets (Mouse/Human, In vitro/In vivo), *TBX6* and *SOX2* are the PreciCE algorithm’s top-ranked transcription factors to perturb for pluripotent-to-mesoderm conversion

Crucially, the algorithm’s output includes both activation and repression. This is key, as activation of some transcription factors may cause broad, unwanted network perturbations. The PreciCE algorithm can compensate for such effects by modulating other genes, thus fine-tuning the network perturbation. As the algorithm searches for an optimal solution to the transcriptional network perturbation problem, it varies λ, a regularization parameter, across a wide range of values. As λ decreases, the predicted perturbation effect changes, increasing the precision of the network perturbation and enabling transcription factor ranking (**Fig. 3F**). The algorithm’s reconstructed network can be visualized, e.g., using pre-existing online tools. This enables users to inspect the effects of predicted perturbations (**Fig. 3G**). Furthermore, we standardized the data pre-processing workflow (**Fig. S8A**). Importantly, to make the model robust and generalizable across contexts, the PreciCE algorithm uses many simplifying assumptions (**Methods**).

We developed a website (https://precice.stanford.edu) where users can run the PreciCE algorithm by drag-and-dropping scRNA-seq datasets, choosing one of our pre-computed gene regulatory networks, and selecting desired starting cell types, target cell types, and undesired competing lineages.

Given that no other algorithms predict the simultaneous up- and downregulation of multiple genes, we first validated our algorithm’s predictions by testing consistency across multiple scRNA-seq datasets. Across three different datasets of pluripotent stem cells (starting state) and mesoderm (target state)^34,35^, and this study **(Methods),** the PreciCE algorithm predicted [*SOX2↓ TBX6↑*] as the most top-ranked perturbation, despite differences in species (mouse/human) and origin (in vitro/in vivo) (**Figs. 3H, S8B-C**). Lower-ranking genes differed. Thus, the PreciCE algorithm showed robustness across datasets but also sensitivity to the nature of the input data.

We next found that the PreciCE algorithm predicted a perturbation for the forced conversion of cardiac fibroblasts to pluripotent stem cells [*SOX2↑ NANOG↑ HAND1↓ HES6↑ POU5F1/OCT4↑*] (**Fig. S8D**). Many genes were plausible based on the literature^36,37^, and we therefore proceeded to test algorithm predictions experimentally.

### The PreciCE algorithm can predict regulators of developmentally specific cell states

The PreciCE algorithm’s consistently high ranking of [*TBX6*↑] for pluripotent-to-mesoderm conversion (**Fig. 3H**) was unexpected. To our knowledge, *TBX6* has only been shown in one study to drive mesoderm differentiation^38^. Thus, a suitable first validation experiment was to see if [*TBX6*↑] could drive mesoderm conversion as efficiently as more well-established mesoderm transcription factors such as *MESP1*, *TBXT* (*T*/*BRACHYURY*), and *MIXL1.* Our computationally reconstructed gene network suggested these and other pro-mesoderm genes would be upregulated as a result of *TBX6* upregulation (**Fig. 3G**).

We performed transient CRISPRa-based activation of *TBX6* or *MESP1* or *TBXT* or *MIXL1* in iPSCs (**Fig. 4A, S9A**, **Methods**) using transfection (using the dCas9/dCas12a system, to make results comparable to later differentiation experiments using this system). As a positive control, we performed a small-molecule-based protocol for cardiomyocyte formation (**Fig. S9B-C)**^39^, collecting cells for scRNA-seq at a time point when mesoderm cells had formed (**Fig, S9D-E**). Three days after transfection of the CRISPR constructs, a fraction of cells had differentiated into cells whose transcriptomes computationally clustered together with those of mesoderm cells from the positive-control condition (**Fig. S9F**) and expressed mesoderm/mesendoderm markers (e.g., *PDGFRA, EOMES, TBXT, MSX1, MESP1, MIXL1*). Interestingly, the [*TBX6*↑] perturbation had generated as many such mesoderm cells as [*MESP1*↑] (17%). This was more than [*TBXT*↑] (5%) or [*MIXL1*↑] (0%) (**Fig. 4B**), indicating that the PreciCE algorithm had accurately predicted *TBX6* as an unconventional mesoderm-inducing transcription factor. The relatively low percentage of differentiation was later improved by enhancing gene modulation efficiency (**Fig. S12A-B**).

**Figure 4.**
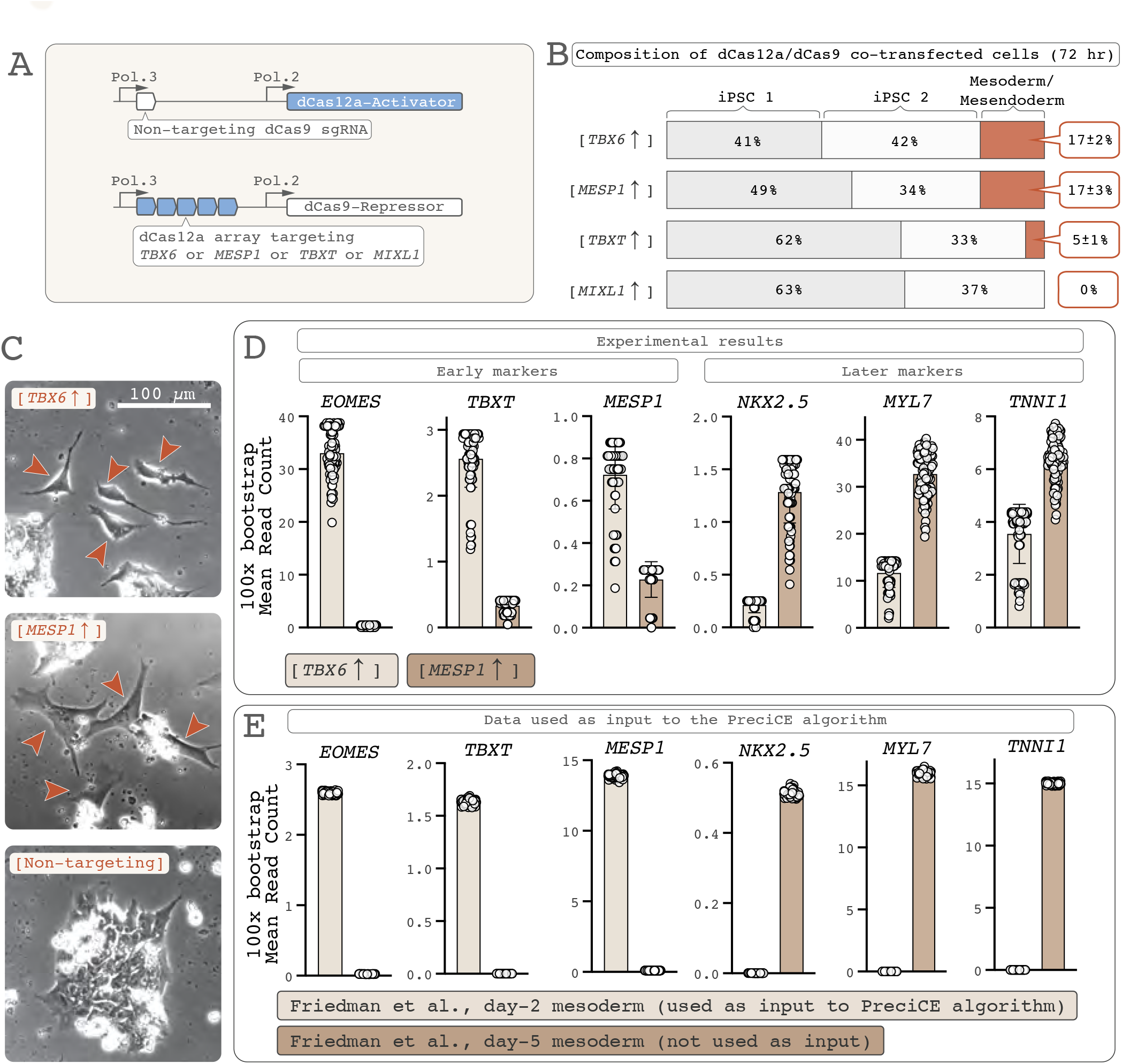
The PreciCE algorithm can predict regulators of developmental stage-specific cell states. (**A**) To validate the PreciCE algorithm’s top-ranked prediction of *TBX6* activation for pluripotent-to-mesoderm conversion, we experimentally compared *TBX6* activation in iPSCs (3 days post-transfection) with other known mesoderm-specific transcription factors using CRISPRa. We used our dCas12a/dCas9 design to make results comparable with later experiments (Fig. 5) and targeted each gene’s promoter with an array of 5 dCas12a gRNAs. (**B**) In the resulting scRNA-seq data, all resulting cells cluster as either *NANOG*(high) iPSCs, *NANOG*(low) iPSCs, or mesoderm/mesendoderm. [*TBX6↑*] and [*MESP1↑*] most efficiently convert iPSCs into mesoderm/mesendoderm. (**C**) Visually, though, [*TBX6↑*] and [*MESP1↑*]-induced cells look different, [*MESP1↑*]-induced cells being larger. (**D**) [*TBX6↑*]-induced cells express markers of earlier mesoderm (*EOMES*, *TBXT/T/BRACHYURY*, *MESP1*) compared to [*MESP1↑*]-induced cells (*NKX2.5*, *MYL7*, *TNNI1*) (Data points represent 100 bootstrapping iterations). (**E**) This makes [*TBX6↑*]-induced cells more similar to the early mesoderm cells that were used as input to the PreciCE algorithm (Friedman et al., small-molecule CHIR99021 protocol, 2-day time point), whereas [*MESP1↑*]-induced cells are more similar to later mesoderm cells, which were not used as input (Friedman et al., 5-day time point). Thus, the PreciCE algorithm accurately predicted a transcription factor (*TBX6*) whose upregulation generated mesoderm of a developmental stage that matched that of the input data.

Interestingly, the cells produced by [*TBX6*↑] and [*MESP1*↑] looked different from one another, the most striking difference being that the [*MESP1*↑]-induced cells were larger (**Fig. 4C**). We found that [*TBX6*↑]-induced cells expressed higher levels of markers associated with early mesoderm/mesendoderm (e.g., *EOMES*, *TBXT*, *MIXL1*, and even *MESP1*) whereas the [*MESP1*↑]-induced cells had higher levels of markers associated with late mesoderm and cardiomyocyte differentiation (e.g., *NKX2.5*, *MYH7*, *MYL7*, *TNNI1*) (**Fig. 4D**). Interestingly, this made the [*TBX6*↑]-induced cells more similar to the cells that had been used as input to the PreciCE algorithm^34^ (**Fig. 4E**). The [*MESP1*↑]-induced cells, in contrast, most resembled later-stage mesoderm found in the same dataset but not used as input to the PreciCE algorithm (**Fig. 4D-E**). This suggested that the PreciCE algorithm predicted an unconventional gene capable of driving the formation of mesoderm matching the developmental stage in the input data (nascent, cardiogenic mesoderm), consistent with a previously described role of *TBX6*^38^.

### Data-driven multi-gene control improves efficiency and precision of cell fate engineering and enables cell fate logic operations

Encouraged by these results, we asked whether PreciCE could improve cell conversion precision as measured by scRNA-seq. We again used the conversion of iPSCs to mesoderm^34^ as our experimental system (**Fig. 3D**). As predicted by the PreciCE algorithm, we performed [*SOX2↓ TBX6↑ NANOG↓ SP5↑*] in iPSCs using our dCas9/dCas12a system (**Fig. S10A**), together with several control perturbations for comparison ([*TBX6*↑], [*TBX6*↑ *SP5*↑], [*SOX2↓ NANOG↓*], [Non-targeting]). Using transfection, gene modulation was transient (CRISPRa: 2-3 days, CRISPRi: ≥5 days, **Fig. S10B-C**). Observing differentiation (**Fig. S10D-E**), we sorted cells for scRNA-seq by Smart-seq3xpress 96 hours post-transfection (without gating for CRISPR construct expression, **Methods**) (**Fig. S10F)**.

We clustered cells from all experimental groups together to gain a global overview of the dataset. At this time point, we observed mesoderm (*MESP1^+^, MSX1^+^*), including both cardiogenic (*TMEM88^+^, HAND1^+^, TNNI1^+^*) and endothelial mesoderm (*PECAM1^+^, KDR^+^*), endoderm (*SOX17^+^, FOXA2^+^, HNF1B^+^*), and two clusters of residual pluripotent stem cells (*NANOG^+^, SOX2^+^, POU5F1^+^*), termed “iPSC 1” (*NANOG*^high^) and “iPSC 2” (*NANOG*^low^, *EOMES^+^*) (**Figs. 5A**, **S10G-H**). Endoderm and endothelial cells had not been present in the PreciCE algorithm’s input data so we classified these and residual stem cells as undesired heterogeneity. Using this dataset, we asked fundamental questions about PreciCE’s performance.

**Figure 5.**
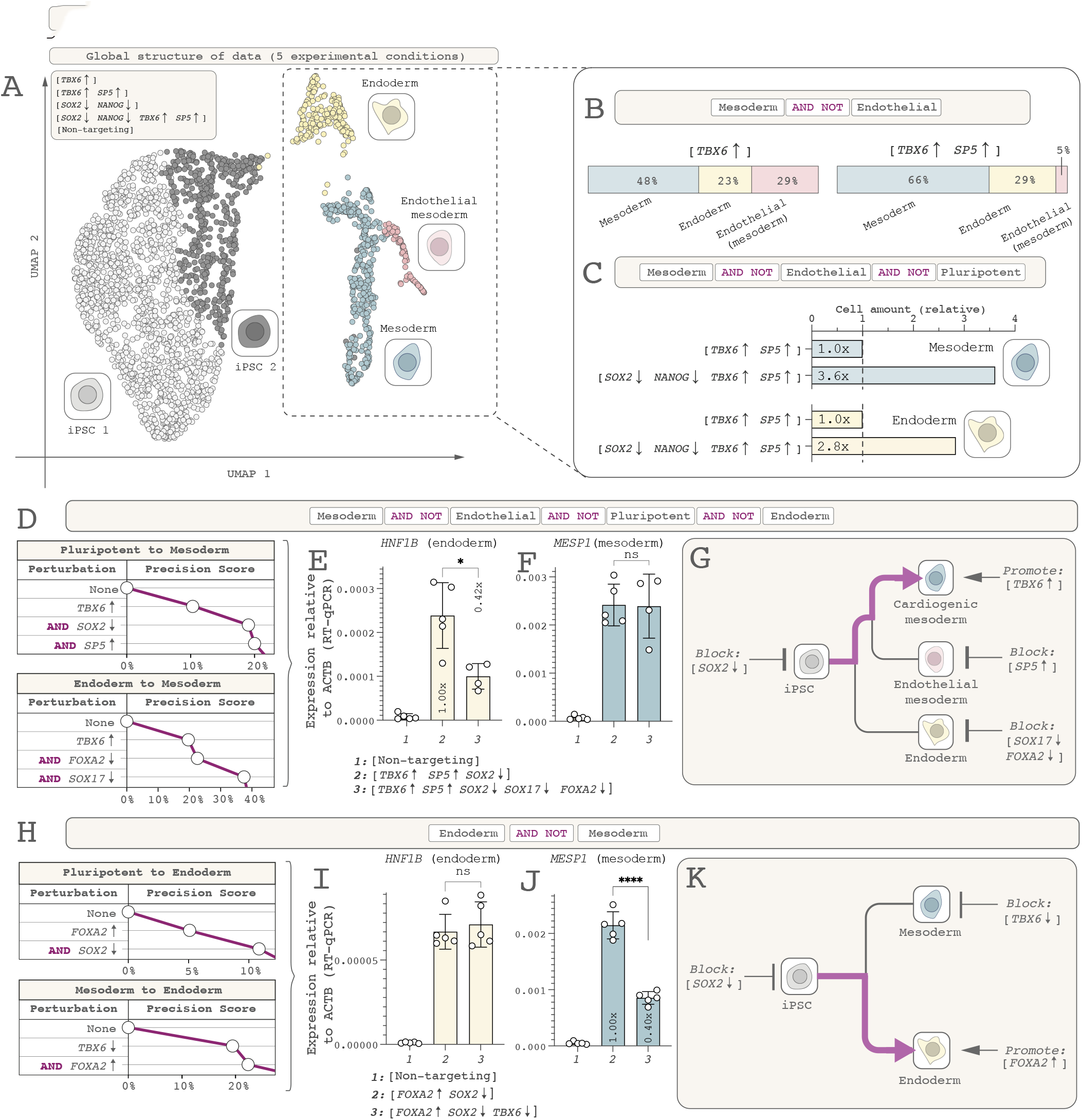
Data-driven precision cell fate conversion using PreciCE. (**A**) For converting iPSCs to cardiac mesoderm, we performed the PreciCE algorithm’s suggested perturbation [*SOX2↓ TBX6↑ NANOG↓ SP5↑*] together with several control perturbations ([*TBX6*↑], [*TBX6*↑ *SP5*↑], [*SOX2↓ NANOG↓*], [Non-targeting]) and FACS-sorted cells for scRNA-seq without gating for transfected cells. We computationally clustered cells from all experimental conditions together. All cells cluster as either undifferentiated iPSCs, cardiogenic mesoderm (desired), endothelial mesoderm (undesired), or endoderm (undesired). The differentiated cells (dashed line) were selected for further analysis: (**B**) Adding *SP5* activation [*TBX6*↑ *SP5*↑] improves specificity by blocking formation of (*SP5*-negative) endothelial cells (Cell-fate logic: *Cardiac mesoderm AND NOT Endothelial*), though not blocking formation of (*SP5*-positive) endoderm. Thus, increasing the number of perturbed genes is one way to rationally include or exclude specific cell fates. (**C**) Adding repression of the pluripotent state [*TBX6*↑ *SP5*↑ *SOX2↓ NANOG↓*] improves efficiency of cell conversion. As expected, it does not markedly affect the mesoderm/endoderm ratio. (Bar chart shows relative number of cells in the scRNA-seq dataset, defined as 1.0x for the [*TBX6*↑ *SP5*↑] condition to facilitate comparison). Thus, destabilizing the starting state is an important feature of PreciCE. (**D**) To actively block endoderm, we input endoderm as “starting state” and mesoderm as “target state” into the PreciCE algorithm, and combined the top-ranked output genes [*TBX6*↑ *FOXA2↓ SOX17↓*] with our pro-mesoderm gene set [*TBX6*↑ *SP5*↑ *SOX2↓*] (excluding NANOG; see main text) to generate the perturbation set [*TBX6*↑ *SP5*↑ *SOX2↓ FOXA2↓ SOX17↓*] for *Cardiac mesoderm AND NOT Endothelial AND NOT Pluripotent AND NOT Endoderm* cell-fate logic. This dramatically reduces formation of *HNF1B*^+^ endoderm (**E**; RT-qPCR data; see **Fig. S9I;** error bars represent standard deviation) while not affecting *MESP1*^+^ mesoderm (**F;** one outlier sample was excluded). (**G**) Thus, data-driven multi-gene modulation can add precision to heterogeneous cell differentiation systems by actively guiding cells through a branched lineage tree. (**H**) The PreciCE algorithm’s predicted gene modulation set for *Endoderm* [*SOX2↓ FOXA2↑*] indeed generates endoderm (**I**) but also some undesired mesoderm (**J**). Combined with an inversion of the Endoderm-to-Mesoderm prediction (**D**) to generate the *Endoderm AND NOT Mesoderm* gene set [*SOX2↓ FOXA2↑ TBX6↓*] actively blocks mesoderm formation (**J**) without affecting endoderm formation (**I-K**), showcasing the modularity and programmability of PreciCE.

First, we asked, is undesired cell fate heterogeneity reduced by increasing the number of perturbed genes? The answer was yes. We compared the [*TBX6*↑] and [*TBX6*↑ *SP5*↑] perturbations (**Fig. 3D**). The addition of [*SP5*↑] almost completely abolished undesired endothelial cell formation (5% vs 29% endothelial cells of all differentiated cells), though undesired endoderm cells were still present (23% vs 29% endoderm cells of all differentiated cells; **Fig. 5B**). This was consistent with *SP5* being expressed by mesoderm and endoderm cells but not by endothelial cells (**Fig. S10J**). In effect, the addition of [*SP5*↑] amounted to a logical A AND B AND NOT C gate, enabling (*Mesoderm) AND (Endoderm) AND NOT (Endothelial)* specification.

Second, we asked, is cell conversion more efficient by actively blocking the starting cell state? Here, too, the answer was yes. We compared the [*TBX6*↑ *SP5*↑] and [*TBX6↑ SP5↑ SOX2↓ NANOG↓*] perturbations and found that the latter improved mesoderm formation 3.6-fold (**Fig. 5C**), effectively executing (*Mesoderm) AND (Endoderm) AND NOT (Endothelial) AND NOT (Pluripotent stem cells)* logic. Yet undesired endoderm cells still remained (**Fig. 5C**).

Third, therefore, we asked, can heterogeneity be further reduced by *actively* blocking closely related but undesired lineages? Here, again, the answer was yes. To actively block endoderm formation, we used the PreciCE algorithm and set endoderm as starting state and mesoderm as target state. We combined this endoderm-to-mesoderm output (**S10L-M**) with our pluripotent-to-mesoderm output to generate the perturbation set [*TBX6↑ SP5↑ SOX2↓ FOXA2↓ SOX17↓*] (**Fig. 5D**). We analyzed the outcome using RT-qPCR against *MESP1* and *HNF1B*, genes almost exclusively expressed in the mesoderm and endoderm lineages, respectively **(Fig. S10N-O**). [*TBX6↑ SP5↑ SOX2↓ FOXA2↓ SOX17↓*] indeed successfully reduced endoderm formation (**Fig. 5E**), without reducing mesoderm formation (**Fig. 5F**), effectively executing (*Mesoderm) AND NOT (Endothelial) AND NOT (Pluripotent stem cells) AND NOT (Endoderm)* logic. This demonstrated that data-driven multi-gene control can enable rational and programmable repression of undesired cell fates (**Fig. 5G**).

To demonstrate the generalizability of PreciCE, we conversely sought to guide cells *toward* endoderm while this time *repressing* mesoderm. Indeed, guided by the PreciCE algorithm (**Fig. 5H, S11A-B**), a perturbation for (*Endoderm) AND NOT (Mesoderm) (*[*FOXA2↑ SOX2↓ TBX6↓*]) converted iPSCs into endoderm while selectively blocking formation of mesoderm (**Figs. 5I-J, S11C-D**). Interestingly, the ability of [*FOXA2↑*] to convert iPSCs into endoderm was completely dependent on the simultaneous repression of *SOX2* (**Fig. S11E**). This was different from iPSC-to-mesoderm conversion, where [*TBX6↑*] had been able to convert iPSCs into mesoderm even in the absence of [*SOX2↓*], albeit inefficiently (**Figs. 4C, 5B, S11F**). This demonstrated the crucial importance of repressing the starting cell state for unlocking the cell programming action of some transcription factors, such as *FOXA2*.

Taken together, these findings demonstrate that data-driven multi-gene control with PreciCE can improve both cell conversion efficiency (by blocking the starting cell state) and reduce heterogeneity (by modulating many genes at once and/or actively blocking competing cell lineages), thus laying the foundation for powerful data-driven cell fate engineering applications.

In parallel, we discovered that cell conversion efficiency could be dramatically enhanced by expressing the Cas genes under the strong CAG promoter instead of the relatively weak hPGK and EFS promoters used for earlier experiments (**Fig. S12A-B**), demonstrating that Cas gene expression levels markedly influence performance of the PreciCE toolbox.

### Engineered iPSC lines enable user-friendly execution of multi-gene modulation

To facilitate use by other researchers, we generated iPSC lines carrying each of the three multi- gene control systems genomically integrated into the *AAVS1* safe-harbor locus and inducible through doxycycline administration (**Fig. S12C-G**). We used one of these lines (**Fig. S12G**) to show that a multi-gene differentiation program (to mesoderm) could be triggered through the simple addition of doxycycline to the cell culture medium (**Fig. S12H**).

## Discussion

We here describe a method for data-driven precision cell fate control and demonstrate that it can be used to improve precision in cell conversion experiments. Specifically, we find that (1) multi- gene perturbation, guided by a uniquely tailored prediction algorithm, can achieve more precise cell states than single-gene perturbation, (2) simultaneous gene activation and repression enhances precision by actively blocking undesired cell states, (3) the seamless interface between computational prediction and experimental multi-gene control enables rational engineering of cell identity.

Our three multi-gene control systems (**Fig. 2**) give users the flexibility to choose the system that best matches their needs. Guide-RNA design and choice are crucial for the success of PreciCE experiments. The number of genes being modulated may dictate use of Pol. III- (≤∼10 gRNAs) or Pol. II-transcribed (>10 gRNAs) single arrays. Desired duration of gene repression may favor mRNA-targeting (Cas13d) or epigenetic (dCas12a, dCas9)-based systems. A desired delivery method may favor a single-array (Cas13d/dCas12a, As-/Lb-dCas12a) or two-construct all-or-none (dCas9/dCas12a) design. And cell type-specific effects^40^ or collateral activity^25^ may restrict a user’s choice of system. Many validated gRNAs and gRNA design tools exist for dCas9, facilitating its use. On the other hand, when no validated gRNAs exist, the ability of dCas12a and Cas13d to encode multiple gRNAs on a single array can make gene modulation more predictable. Some of the systems’ features are interchangeable. For example, a two-construct all-or-none architecture can be used for the Cas13d/dCas12a hybrid system. And a dCas12a activator can be replaced with a dCas9 activator. Importantly, the use of engineered hybrid CRISPR arrays greatly simplifies the engineering of multi-gene networks, enabling the delivery of multi-gene programs as single data packets. This can reduce heterogeneity caused by variations in gRNA delivery and expression efficiency, as often seen with previous dCas9-based methods.

Recent advances on defining novel transcriptional and epigenetic effectors provide a reservoir of useful molecules that can be modularly integrated into the PreciCE platform. For example, researchers can use novel epigenetic activators (e.g., TET1^41^, p300^42^, NFZ^43^) or repressors (DNMT^44^, ZIM3-KRAB^45^, etc.) for simultaneous activation and repression. These effectors can perform new modes of regulation (e.g., long-duration silencing), thus expanding the functionality of PreciCE.

The success of PreciCE relies on using efficient gRNAs. Combining multiple non-optimal gRNAs into a single array could fail to cause any gene modulation. In this regard, one major benefit of the single-array-based gRNA expression is the ability to encode multiple gRNAs targeting the same gene on a single array, thus circumventing time-consuming gRNA optimization. Furthermore, the single-array system has great potential for large-scale multiplexed screening. Pooled arrays can be synthesized, each encoding desired activating guides and repressive guides in pre-defined stoichiometry, enabling elucidation of genetic networks in novel ways.

Natural differentiation often, though not always^46^, proceeds through a series of transient intermediate states. Currently, the PreciCE algorithm is only fed information about starting and target cell states. This may impact the ability to execute conversions to developmentally distant target states that may involve multiple intermediate stages. However, combining PreciCE with other cell differentiation protocols and activating the PreciCE algorithm’s predicted gene set during crucial moments in a cell’s differentiation trajectory may enhance precision with current protocols.

The PreciCE algorithm requires using pure starting and target populations as input. If the input is a heterogeneous mix of cell types (e.g., clustering together in a UMAP), the algorithm’s prediction will be based on that mix. One reason we observed undesired endoderm cells in our mesoderm conversion was likely that the input data expressed not only mesoderm markers but also earlier mesendoderm markers. Furthermore, the algorithm is sensitive to noise, especially caused by very small datasets. This manifests in the form of networks with noisy or incorrect edges as well as genes that may get overlooked in the differential expression due to batch effects or other experimental artifacts. In our experience, the starting and target cell types should each contain at least several hundred cells.

Collectively, these findings establish PreciCE as a powerful platform for data-driven cell fate engineering. By leveraging growing databases from cell atlases, PreciCE could become an essential asset in the generation of human cells on demand and help transform the manipulation of cell identities into a rational and programmable discipline.

## Author contributions

J.P.M. and S.L.Q. conceived the study. J.P.M. performed experiments and data analysis, supervised experiments, wrote the manuscript, and made figures. Y.R. developed the PreciCE algorithm. D.S. developed the PreciCE website, performed statistical analyses for multi-gene regulation in single cells, and performed experiments. Y.S. created the stably integrated iPSC lines. P.R.C.T performed experiments. R.S. supervised scRNA-seq experiments and transcriptional bursting analyses. J.L. supervised Y.R. L.S.Q. supervised the project.

## Acknowledgments

We thank Michael Hagemann-Jensen and Christoph Ziegenhain (Karolinska Institutet) for their great help with the Smart-seq3xpress workflow. We thank Matthew Lau (Stanford University) for help with the PreciCE website, and Timothy Abbott (Stanford University) for the generation of the GFP-expressing HEK293T cell line. This work was funded by a California Institute for Regenerative Medicine (CIRM) DISC2 Award (#DISC2-12669), Chan Zuckerberg Neurodegeneration Challenge Network NDCN (#DAF2022-250596), and NIH National Cancer Institute R01 (#R01CA266470). J.P.M. was supported by the Human Frontier Science Program Long-Term Fellowship (LT000955/2018), David och Astrid Hageléns Stiftelse, Magnus Bergvalls Stiftelse, and the Sweden-America Foundation. R.S. was supported by the Swedish Research Council (no. 2017-01062) and the Torsten Söderberg Foundation. J.L. was supported by NSF under Nos. OAC-1835598 (CINES), CCF-1918940 (Expeditions), DMS-2327709 (IHBEM); Stanford Data Applications Initiative, Wu Tsai Neurosciences Institute, Stanford Institute for Human-Centered AI, Chan Zuckerberg Initiative, Amazon, Genentech, GSK, Hitachi, SAP, and UCB. L.S.Q. acknowledges support by NSF CAREER award 2046650 (L.S.Q.), NIH Director’s Pioneer Award DP1NS137219 (L.S.Q.), and CHAU HOI SHUEN FOUNDATION. L.S.Q. is a Chan Zuckerberg Biohub investigator.

## Data availability

ScRNA-seq data will be deposited in a public repository.

## Availability of cell lines and plasmids

Cell lines and plasmids will be deposited in public repositories.

## Computer code availability

The code for the PreciCE algorithm is available at https://github.com/snap-stanford/precice. The PreciCE Online Tool is available at https://precice.stanford.edu.

## Methods

### CRISPR-based multi-gene control experiments

#### dCas12a gene

We used nuclease-deactivated hyper-Cas12a derived from *Lachnospiraceae bacterium*^21^ followed by an activator or repressor, and GFP or mCherry. For the Lb-dCas12a/As- dCas12a system, we used enhanced-dCas12a from *Acidaminococcus species*^48^.

#### Cas13d gene

We used *Ruminococcus flavefaciens* Cas13d^22^.

#### dCas9 gene

We used nuclease-deactivated *Streptococcus pyogenes* Cas9^49^ fused to a repressor domain and mCherry.

#### HEK293T cells

For development of the CRISPR-based multi-gene control systems (**Fig. 2**), we used HEK293T cells from Takara Bio (Japan) carrying (1) a lentivirally integrated GFP tagged to a PEST degradation domain (enabling rapid protein dynamics suitable for gene repression experiments) expressed under a TRE3G promoter, and (2) a reverse tetracycline transactivator (Rtta) expressed under the strong constitutive EF1ɑ promoter, enabling rapid GFP expression through addition of doxycycline to the culture medium. These cells showed low but detectable GFP expression even in the absence of doxycycline, enabling us to test Cas13d-based gene repression without inducing collateral activity.

#### iPSCs

We used iPSC line SCVI-480, kindly provided by Joseph C. Wu, MD, PhD at the Stanford Cardiovascular Institute funded by NHLBI BhiPSC-CVD 75N9202D00019. These cells were derived from blood cells drawn from a healthy, 18-year-old woman of African American descent, with consent for research use. All experimental procedures were approved by Stanford’s Stem Cell Research Oversight (SCRO) panel.

#### Cell culture of HEK293T cells

Cells were grown in DMEM with high glucose (4.5 g/L), GlutaMAX, and pyruvate (Gibco; 10569-010) supplemented with 10% fetal bovine serum (Sigma-Aldrich, St. Louis, MO, USA; F7524-500ML) and 1% penicillin-streptomycin (Gibco; 15070063). For passaging, cells were dissociated with TrypLE (Thermo Fisher; 12604039).

#### Cell culture of hiPSCs

Cells were grown in 6-well plates coated with Laminin-521 (BioLamina, Sundbyberg, Sweden; 17187501) according to BioLamina’s protocol (1/10 dilution in PBS for 2 hr). Cells were kept in mTeSR Plus (Stemcell Technologies; 17187501), which was changed daily. For passaging, cells were dissociated with Accutase (Corning Inc., Corning, NY, USA; 15323609) followed by addition of 1 volume of mTeSR Plus containing 1x RevitaCell Supplement (Gibco; A2644501). Cells were transferred to a 15 ml Falcon tube and centrifuged at 300*g for 4 minutes. Thereafter, supernatant was removed, 2 ml new mTeSR Plus containing RevitaCell Supplement was added. Cells were counted using a Countess 3 Automated Cell Counter (Thermo Fisher) by performing four counts on the same chip and using the average cell number to increase seeding precision. We seeded 150,000 or 300,000 cells per well in a 6-well plate in mTeSR Plus containing RevitaCell Supplement. For subsequent medium changes, RevitaCell Supplement was excluded. Cell cryopreservation was performed using BamBanker (Fujifilm Wako, Neuss, Germany; 306-14684).

#### Cell differentiation using a small-molecule (CHIR99021) protocol

For mesoderm differentiation using small molecule CHIR99021, we adapted a previously published protocol^57^. Briefly, wells in a 24-well plate were coated with Laminin-521 (BioLamina, LN521-02; Thermo Fisher, #A29249) at 1/10 dilution in PBS for 2 hr at 37°C. iPSCs were seeded at a density of 100,000 cells per well in mTeSR Plus (#100-0274 containing supplement #100-0275, STEMCELL Technologies, BC, Canada), containing 1x RevitaCell Supplement (#A2644501, Thermo Fisher). Twenty-four hr after seeding, medium was changed to mTeSR Plus without RevitaCell Supplement. Forty-eight hr after seeding, medium was changed to RPMI supplemented with L-glutamine and HEPES (#22400089, Gibco, MT, USA) and 1x B27-minus-insulin (#A1895601, Gibco) and containing 10 µm CHIR99021 (#SML1046, Sigma-Aldrich, MO, USA). Exactly 72 hr after seeding, medium was changed to RPMI (with L-glutamine and HEPES) containing 1x B27- minus-insulin, but without CHIR99021. Ninety-six hr after seeding, cells were dissociated and FACS-sorted for scRNA-seq.

#### Cell differentiation using CRISPRa/CRISPRi

For mesoderm differentiation using single-gene upregulation (**Fig. 4)**, we used the dCas9/dCas12a system (**Fig. 2G**; “Experiment #23-006”) to make results comparable to subsequent differentiation experiments, even though no dCas9- based gene repression was performed in this experiment. We targeted each gene with 5 dCas12a gRNAs in a single array. We co-transfected iPSCs with the dCas12a-Activator and dCas9- Repressor constructs using reverse transfection. Twenty-four hr after transfection, medium was changed to RPMI supplemented with L-glutamine and HEPES (#22400089, Gibco, MT, USA) and 1x B27-minus-insulin (#A1895601, Gibco). Forty-eight hr after transfection, medium was again changed to RPMI containing B27-minus-insulin. Seventy-two hr after transfection (a time point where pilot experiments showed *MESP1* upregulation and morphological changes, both suggestive of mesoderm differentiation), cells were dissociated and resuspended in PBS containing RevitaCell Supplement and FACS-sorted into 384-well plates for scRNA-seq (see below), gating for GFP and mCherry present on the dCas12a-Activator and dCas9-Repressor constructs, respectively.

For mesoderm and endoderm differentiation using multi-gene up- and downregulation (Fig. 5; “Experiment #23-001”), we used the dCas9/dCas12a system (**Fig. 2G**) similar to the setup in the previous paragraph. But instead of changing to RPMI + B27-minus-insulin medium already 24 hr post-transfection, we instead changed to mTeSR Plus (without RevitaCell Supplement) 24 hr post-transfection. At 48 and 72 hr post-transfection, we changed medium to RPMI + B27-minus-insulin. At 96 hr post-transfection (a time point where pilot experiments showed *MESP1* upregulation and morphological changes (both suggestive of mesoderm differentiation), cells were analyzed, either by dissociation and FACS-sorting for scRNA-seq, or by lysis and downstream processing for RT-qPCR. For Fig. 5A-C, we sorted cells without gating for the fluorescent proteins fused to the Cas protein, as these protein constructs had largely been diluted out by cell division at this time point.

#### Generation of scRNA-seq libraries

We generated scRNA-seq libraries using the Smart- seq3xpress method and sequenced on a DNBSeq-G400RS platform (MGI, Shenzhen, China), as described in ref.^26^, using either HotMPS or StandardMPS chemistry. Smart-seq3xpress involves cell sorting by FACS into 384-well plates and downstream processing to achieve full-transcript coverage. We generated scRNA-seq datasets from the following experiments, using the following specifications:

***(1) Cas13d/dCas12a hybrid array testing in HEK293T cells*** (**Figs. 2C, S3);** “Experiment #22-001”): Forty-eight hr post-transfection, cells were dissociated and stained with an A647- conjugated antibody against the target gene product CD9 (#341648, BD Biosciences, Franklin Lakes, NJ, USA). Cells were suspended in PBS and FACS sorting was done using a BD Influx (BD Biosciences, Franklin Lakes, NJ) using a 140-um nozzle. Cells were sorted for uptake of the CRISPR array plasmid (BFP) and dCas12a-miniVPR and Cas13d plasmids (mCherry) and CD9- A647^high^ expression. During library preparation, we performed 12 cycles of pre-amplification PCR and 12 cycles of index PCR and used 0.002 ul TDE1 Tn5 enzyme per cell (Illumina Tagment DNA TDE1 Enzyme and Buffer Kits; Illumina, San Diego, CA, USA). Sequencing was performed using an SE100 sequencing kit (MGI). After sequencing and quality-control, the dataset consisted of 688 cells.
***(1) Lb-dCas12a/As-dCas12a hybrid array testing in HEK293T cells*** (**Fig. 2F**; “Experiment #24-019”): Forty-eight hr post-transfection, cells were dissociated and suspended in PBS containing a live/dead marker (NucRed Dead 647; Thermo Fisher). FACS sorting was done using a BD FACS Melody (BD Biosciences) using a 100-um nozzle. Cells were sorted based on Lb- dCas12a-mCherry-KRAB expression and low fluorescence of the Live/Dead marker. During library preparation, we performed 12 cycles of pre-amplification PCR and 12 cycles of index PCR and used 0.003 ul TDE1 Tn5 enzyme per cell. Sequencing was performed using a PE100 sequencing kit (MGI). After sequencing and quality-control, the dataset consisted of 1334 cells. However, because our BD FACS Melody was not equipped with a UV laser, we had been unable to sort for cells expressing the As-dCas12a-VPR-BFP construct. We therefore subsetted the dataset to include only cells expressing high levels of the As-dCas12a-VPR-BFP (>100 reads per cell), yielding a final dataset of 543 cells.
***(1) dCas9/dCas12a all-or-none architecture testing in HEK293T cells*** (**Fig. 2I**; “Experiment #24-022”): Forty-eight hr post-transfection, cells were dissociated and suspended in PBS. FACS sorting was done using a BD FACS Melody (BD Biosciences) using a 100-um nozzle. Cells were sorted based on dCas9-mCherry-KRAB expression and dCas12a-miniVPR-GFP expression. During library preparation, we performed 12 cycles of pre-amplification PCR and 12 cycles of index PCR and used 0.004 ul TDE1 Tn5 enzyme per cell. Sequencing was performed using a PE100 sequencing kit (MGI). After sequencing and quality-control, the dataset consisted of 1003 cells.
***(1) iPSC-to-mesoderm differentiation using CHIR99021*** (**Figs. 3H, 4, S9** “Experiment #24- 010”): See experimental settings in the *Small-molecule mesoderm differentiation* section. At the end of the experiment (96 hr post-seeding), cells were dissociated and suspended in PBS containing DAPI (4 µg/ml) as a live/dead marker. FACS sorting was done using a BD Influx (BD Biosciences) using a 140-um nozzle. During library preparation, we performed 12 cycles of pre-amplification PCR and 12 cycles of index PCR and used 0.003 ul TDE1 Tn5 enzyme per cell. Sequencing was performed using a PE100 sequencing kit (MGI). After sequencing and quality-control, the dataset consisted of 476 cells.
***(1) iPSC differentiation using [TBX6↑] and [MESP1↑] and [MIXL1↑] and [TBXT↑]*** (**Figs. 4, S8**; “Experiment #23-006”): See experimental settings in the *Differentiation using CRISPRa/CRISPRi* section. At the end of the experiment (72 hr post-transfection), cells were dissociated and suspended in PBS containing DAPI (4 µg/ml) as a live/dead marker. FACS sorting was done using a BD Influx (BD Biosciences) using a 140-um nozzle, gating for dCas9-mCherry-KRAB expression and dCas12a-miniVPR-GFP expression. During library preparation, we performed 12 cycles of pre-amplification PCR and 12 cycles of index PCR and used 0.003 ul TDE1 Tn5 enzyme per cell. Sequencing was performed using a PE100 sequencing kit (MGI). After sequencing and quality-control, the dataset consisted of 1179 cells. We further computationally subsetted the dataset by including only cells that contained ≥1 read of any transfected construct, yielding a final dataset of 717 cells.
**(1) *iPSC differentiation using [TBX6↑] and [TBX6↑ SP5↑] and [SOX2↓ NANOG↓ TBX6↑ SP5↑] and [SOX2↓ NANOG↓]*** (**Figs. 5A-C, S10**; “Experiment #23-001”): See experimental settings in the *Differentiation using CRISPRa/CRISPRi* section. At the end of the experiment (96 hr post-transfection), cells were dissociated and suspended in PBS containing DAPI (4 µg/ml) as a live/dead marker. FACS sorting was done using a BD Influx (BD Biosciences), using a 140-um nozzle. Because transfected plasmids have largely been diluted out at this time point, we did not gate for expression of dCas9-mCherry-KRAB or dCas12a-miniVPR-GFP. This meant that many sorted cells were untransfected. During library preparation, we performed 12 cycles of pre-amplification PCR and 12 cycles of index PCR and used 0.002 ul TDE1 Tn5 enzyme per cell. Sequencing was performed using a SE100 sequencing kit (MGI). After sequencing and quality-control, the dataset consisted of 2959 cells.

#### Flow cytometry

We performed flow cytometry and sorting using BD FACSMelody (100-µm nozzle) or BD Influx (140-µm nozzle) or BD FACSAria Fusion (100-140-µm nozzle) (BD Biosciences). We also used CytoFLEX (Beckman Coulter, Brea, CA) for analysis. FlowJo v.10.7.1 was used for data processing.

#### RT-qPCR

When collecting cells for RT-qPCR, cells were dissociated in bulk and were thus not sorted based on uptake of CRISPR constructs. An exception was the Cas13d/dCas12a system (**Fig. S2U**), where samples were FACS-sorted into lysis buffer. RNA was extracted using the RNeasy Plus Mini Kit (Qiagen, Germany), as recommended by the manufacturer. gDNA removal was performed using TURBO DNA-free kit (Thermo Fisher) after RNA extraction. Reverse transcription was performed using Maxima H Minus Reverse Transcriptase (Thermo Fisher). Quantitative PCR reactions were run on a ViiaA 7 Real-Time PCR System (Applied Biosystems, MA, USA) using Power SYBR Green PCR Master Mix (Thermo Fisher). ACTB was used as reference gene. All expression values were calculated using each sample’s own ACTB value as reference, and “expression relative to ACTB” (2^-dCt^) was used for all plots.

### scRNA-seq computational analysis

#### Primary data processing

zUMIs^58^ (v.2.8.2 or newer) was used to process raw FASTQ files as described in ref.^26^.

#### Quality control

After sequencing and read mapping, a reads-per-cell versus percent-exonic- reads plot informed exclusion of failed libraries.

#### Differential expression

For calculating CRISPRa/i-based expression fold-change for scRNA-seq data (Fig. 2C, F, I), we compared pseudobulk expression level in the following way. We summed all sequencing reads for all cells expressing the targeting gRNA and divided this by the corresponding number for cells expressing non-targeting gRNAs. We chose this method as it best approximates how RT-qPCR-based expression fold-change is measured.

#### Heatmaps

**(Fig. 2C, F, I)**. Heatmaps were plotted in R using the Heatmaply package (v.1.5.0)^59^, using data matrices containing all reads (as opposed to UMI-only). Note that cells were not hierarchically clustered but listed in arbitrary (alphabetical) order.

#### Violin plots

(Figs. S3C, S4J, S5G). We plotted violin plots in R using the ggplot2 package (v.3.5.0)^68^. Plots were made using expression data of all reads (as opposed to only UMI reads).

#### UMAP and dataset integration

UMAPs were produced using Seurat v. 5.0.1^60^ using expression matrices of all reads or UMI reads and the 2000 most variable genes. For integrating the two datasets in **Fig. S9F** (small-molecule-mediated differentiation and CRISPRa-mediated differentiation) using Seurat, we split the datasets by experimental condition and integrated using the JointPCAIntegration method.

#### Volcano plots

(Figs. S3E, S4K, S5H). We performed differential expression analysis and plotted volcano plots in R using edgeR^61^. We arbitrarily set 1.5 as an expression fold-change cutoff for significantly expressed genes. Note that fold-change values for some genes differ dramatically from those in **Fig. 2C, F, I**, as Limma/Voom calculates fold-change differently when one value is zero or near-zero.

#### Analysis of how many genes were perturbed in individual cells (Fig. S6)

A scRNA-seq expression matrix was loaded into Python using Scanpy (v.1.10.2). Statistical analyses were performed using SciPy (v.1.14.0).

First, we estimated how many genes were perturbed in individual cells (**Fig. S6A-C**). Conceptually, this analysis was done in the following way. For each of the 4-6 CRISPR target genes, we first looked at that gene’s “baseline” expression level in control cells expressing the non-targeting CRISPR array. That baseline level corresponded to a distribution of sequencing reads (**e.g., Fig. S3C**). We used this baseline expression distribution to set an arbitrary threshold, beyond which that gene would be considered “perturbed”. Having set such thresholds for all 4-6 target genes, we then looked at each cell expressing the *targeting* CRISPR array (our experimental group). For each such cell, we asked if the first target gene was classified as perturbed. We next asked if that same cell had the second target gene perturbed, and the third, and so on. We repeated this procedure for all cells. Then, the number of genes classified as perturbed in each individual cell was plotted in **Fig. S6A-C**. Technically, the analysis was done using a one-sided Wilcoxon signed-rank test, generating p-values to assess statistical significance. We used the Benjamini–Hochberg procedure to create the significance threshold for determining if each gene was perturbed in any given cell. We performed bootstrapping by excluding 20% of samples per iteration randomly and sampling with replacement over 100 iterations. Visualizations were performed using the Seaborn and Matplotlib libraries.

Note that on the scRNA-seq level, gene activation and repression do not generate binary on-off expression patterns but rather shifts in sequencing read distributions (**Figs. S3C, S4J, S5G**). Therefore, this single-cell-level analysis has noise. For example, because of the sharp thresholds for when a gene would be considered perturbed, some control cells (expressing the non-targeting array) were also classified as perturbed for some genes (**Fig. S6A-C**). And because of zero-inflation in scRNA-seq data, the number of targeting cells showing multi-gene perturbation is likely an underestimate.

To analyze whether each individual cell showed some bias for or against regulating all 4-6 target genes (**Fig. S6D-F**), we performed the following steps. First, given the thresholds described above (beyond which a target gene was considered perturbed), we analyzed how many percent of cells were classified as “perturbed” for each target gene. If target genes are independently modulated, the fraction of cells showing *all* genes perturbed should be equal to the multiplication product of these percentages. In the same way, we calculated the expected fraction of cells showing subsets of genes perturbed **(Fig. S6D-F)**. We compared these expected fractions with the actual observed fractions to robustly and confidently analyze whether cells showed biases for or against multi-gene modulation on the single-cell level.

#### Transcriptional bursting analysis

This analysis was performed as described previously ^33^.

#### The PreciCE Algorithm

Our goal was to design a computational model that leverages single-cell RNA-seq data from source and target cell populations to predict the most effective genes to perturb, identifying the 𝑘 best genes for a successful transition where 𝑘 is a variable integer. Additionally, this model evaluates the efficiency of various perturbation combinations, facilitating the optimal selection of 𝑘 based on resource constraints.

Let 𝑔_𝑖_ be a gene whose expression is measured using single-cell RNA sequencing technology. Let 𝐺 be the set of all such genes and let 𝑇 ⊂ 𝐺 be the set of all transcription factors. Let 𝒙 ∈ ℝ^|𝐺|^ be the vector of expression values for all genes in any given cell. We define 𝜃 ∈ ℝ^|𝐺|^to be a perturbation to this expression vector containing at most |𝑇| non-zero values. The impact of this perturbation on the complete transcriptional state can be represented as 𝒙_𝜃_, using a gene regulatory model 𝑓 that maps from ℝ^|𝐺|^ → ℝ^|𝐺|^.

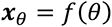

Let 𝒙_𝒔_, 𝒙_𝒕_ ∈ ℝ^|𝐺|^ represent two distinct transcriptional states (the terms ’gene expression vector’ and ’transcriptional state’ are used interchangeably). Our goal is to transform the cell from state 𝑥_𝑠_ to 𝑥_𝑡_ using a perturbation 𝜃^3^ that consists of only 𝑘 ≤ |𝑇| non-zero entries. This can be represented as the following optimization problem:

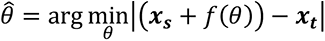

##### Learning 𝑓

Let 𝑇 ⊂ 𝐺 be the set of all genes that code for transcription factors. We define 𝒗 ∈ ℝ^|𝐺|^ to be the expression values of transcription factors alone and zero elsewhere, that is 𝑣_𝑖_ = 𝑥_𝑖_ ⋅ 𝟙[𝑔_𝑖_ ∈ 𝑇] for all 𝑣_𝑖_ ∈ 𝒗, 𝑥_𝑖_ ∈ 𝒙. We also define the notation 𝒗^(–j)^ which is equivalent◻ to 𝒗, except that 𝑣_j_ has been set to zero. 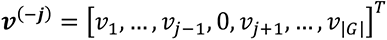. Lastly, we define the gene regulatory model for a specific target gene 𝑔_j_ as 𝑓_j_ from ℝ^|𝐺|^ → ℝ. This elementwise formulation is useful for learning non-linear functions.

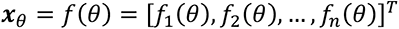

##### Subproblems

This larger problem is composed of the following subproblems:

*1. Inferring an unweighted network of interactions between transcription factors and genes:* Determine the matrix 𝐄 ∈ {0,1}^|𝐺|×|𝐺|^ such that ∀𝑒_𝑖j_ ∈ 𝐄, 𝑒_𝑖j_ = 1 if gene 𝑔_𝑖_ has an impact on the expression of gene 𝑔_j_. In the simplest case, 𝑓 = 𝐄 ⋅ 𝜃 and the second step is not performed.
*2. Determining direct transcriptional relationships between transcription factors and genes:* Given 𝐄, determine 𝑓 (as defined previously).
*3. Identifying the minimal set of transcription factors to perturb to achieve the desired target state:* Identify a sparse transcription factor perturbation vector 𝜃 such that ;<𝒙_𝑠_ + 𝑓(𝜃)> − 𝒙_𝑡_; is minimized.

While 1 and 2 seem closely related, they have historically been approached as separate problems with separate validation sets. Hence, we partition them as well.

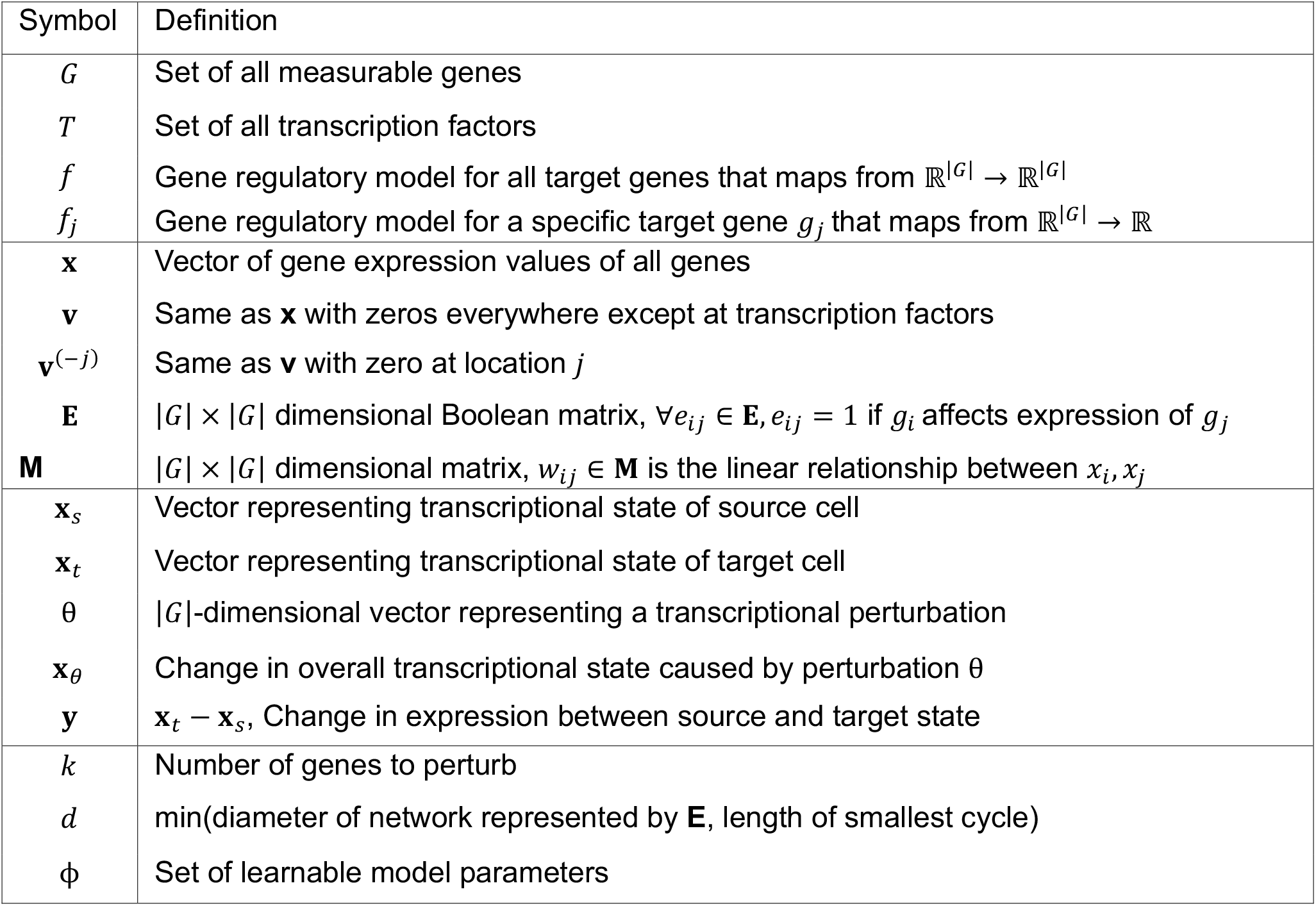

##### Approach

We describe our methodology for the subproblems identified in the previous section.

##### Inferring an unweighted network of interactions between transcription factors and genes

We use the GENIE3 package^62^ for this subproblem, specifically the GRNBoost2 implementation^63^, which has shown superior performance in the accuracy and consistency of detected gene regulatory network edges.

GENIE3 breaks down the problem of recovering the underlying regulatory network into multiple subproblems of identifying regulators for particular genes. However, it does not define the direct transcription factor-gene transcriptional relationship (𝑓). It only creates a list of transcription factor and target gene pairs where each transcription factor has a significant impact on target gene expression (we capture this as 𝐸). By using an ensemble of regression trees, the model recursively splits the training data on each feature while maximizing entropy. The output is a ranking of features (transcription factors) by their importance for predicting the expression of the target gene. While the relationship 𝑓 between the transcription factors and the target genes could be inferred from this model, it’s usually a non-smooth function that would be difficult to optimize over for part (3).

The SCENIC package^64^ further augmented the output from GENIE3 by filtering out interactions where the transcription factor binding motif is not significantly enriched upstream of the target gene sequence^64^. This ensures that each transcription factor is only connected to downstream genes that it directly interacts with and not those genes that may be indirectly impacted. We use this filtering to remove indirect or potentially spurious interactions between transcription factors and downstream genes and consider only those relationships that are validated by sequence information.

##### Learning transcriptional relationships between transcription factors and genes

In the following models, 𝜃 is a |𝐺|-dimensional vector representing any transcriptional perturbation and 𝑥_𝜃_ is the resulting change in overall transcriptional state caused by 𝜃.

*Binary model:* As defined previously, let **E** correspond to a Boolean matrix of all non-zero relationships between transcription factors and target genes (as determined by GENIE3). To prioritize interactions that are within a closer hop distance, we divide the magnitude of the perturbation at each hop by the number of hops so far (ℎ). This is similar to the procedure followed by ref. ^13^. Let 𝑑 = min(diameter of the network defined by **E**, number of edges in the shortest cycle). We do not include self-edges.

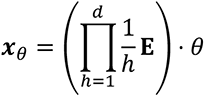

*Linear model:* Let 𝐌 ∈ ℝ^|𝐺|×|𝐺|^ represent an adjacency matrix defining relationships between nodes 𝐺. Let 𝑤_𝑖j_ ∈ 𝐌 represent the weight of an edge between gene 𝑔_𝑖_ and gene 𝑔_j_.

##### Training function

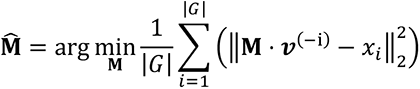

##### Inference function

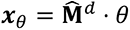

##### Multilayer perceptron

Let 𝑓_𝜙_ represent a multilayer perceptron parametrized by the set of learnable parameters 𝜙. Let 𝑀_1_ ∈ ℝ^𝑛1×|𝐺|^, 𝑀_2_ ∈ ℝ^𝑛2×𝑛1^ , 𝑀_3_ ∈ ℝ^1×𝑛2^ be weight matrices, where 𝑛_1_, 𝑛_2_ are tuned hyperparameters (Setting for current results: 𝑛_1_ = 20, 𝑛_2_ = 10). ReLU represents the elementwise ReLU operator.

##### Training function

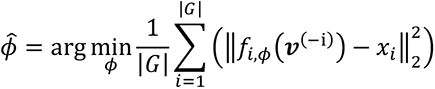

where

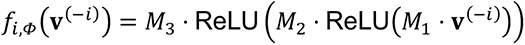

##### Inference function

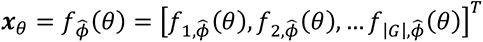

We compared the gene expression predictive performance of the different model formulations for 𝑓 (Table 1), and chose to proceed with the linear formulation.

**Table 1:** Results for predicting target gene expression (𝑓). Average performance across models trained on 9000 random target genes (tasks).

| Train Dataset | Test Dataset | # of cells | Model | RMSE | R<sup>2</sup> |
| --- | --- | --- | --- | --- | --- |
| Duan et al. (Heart) | Duan et al. | 1764 | Binary | 0.018 | -0.296 |
| Duan et al.(Heart) | Duan et al. | 1764 | Linear | **0.010** | **0.491** |
| Duan et al. (Heart) | Duan et al. | 1764 | MLP | 0.023 | 0.326 |
| Duan et al. + Tabula Muris (Heart) | Duan et al. | 2417 | Linear | **0.010** | **0.453** |
| Duan et al. + Tabula Muris (Heart) | Duan et al. | 2417 | MLP | 0.013 | 0.410 |
| Duan et al. + Tabula Muris (5 organs) | Duan et al. | 23433 | Linear | **0.011** | **0.434** |
| Duan et al. + Tabula Muris (5 organs) | Duan et al. | 23433 | MLP | 0.014 | 0.416 |

##### Identifying minimal set of transcription factors to perturb to achieve desired target state

We set the difference in transcriptional state between the source (𝐱_𝑠_) and the target (𝐱_𝑡_) cell type to be 𝒚, where 𝒚 = |𝒙_𝑡_ − 𝒙_𝑠_|.

Our goal then is to identify 𝜃^3^ that minimizes the absolute difference ;𝒙_𝒕_ − <𝒙_𝒔_ + 𝑓(𝜃)>; = |𝑓(𝜃) − 𝒚|. The solution must also satisfy the constraint that only 𝑘 genes must be non-zero, which is incorporated using 𝐿_1_ regularization for enforcing sparsity in 𝜃.

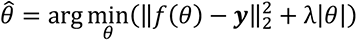

##### Special case: Linear model

In the case where 𝑓 is a linear function and it represents a matrix 𝐌^𝑘^, we can identify 𝜃 using regularized least square regression (lasso) by solving the following optimization problem.

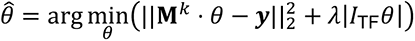

Here 𝐼_TF_ refers to the identity matrix with ones only at the indices corresponding to transcription factors, since those are the only values set by the model. We solve this least-squares regression problem using coordinate descent. The value of 𝜆 is varied over a large range and solutions with desired sparsity are returned.

### Preprocessing single-cell gene expression data

We normalized gene expression in each cell by total counts over all genes. This was followed by a log transformation. We used highly variable genes for the analysis with mean log transformed expression values of greater than 0.0125 and dispersion of 0.5. Additionally, we removed mitochondrial and ribosomal genes from the analysis.

### Computing differential expression (y)

The PrecICE algorithm is very sensitive to the measured differential expression between source and target cell state. Thus, identification of the source and target cell state clusters can have a large impact on the predicted transcription factors to perturb.

- In cases where batch-level variation was present, we used scVI^65^ to remove these effects as well as to measure differential expression between source and target cell state. We used a cutoff of statistical significance (p=0.02) and not the magnitude of differential expression.^66,67^
- In cases where source and target cell state were measured as part of the same experiment without large sources of variation, we did not perform batch correction. For computing differential expression between source and target cell states, we used Seurat with default parameters. We used a cutoff of statistical significance (p=10^-6^) and not the magnitude of differential expression.^67^

### The Precision Score

As part of its output, the PreciCE algorithm produces a Precision Score (**e.g.,** **Fig. 3D**). The Precision Score is equal to the predicted ‘error’ reduction between the desired target cell state and the state achieved by the suggested perturbation (as predicted by the model). This value is normalized by the total error between the original source state and the target state (|𝑓(𝜃) − 𝑦|)/𝑦. The Precision Score can thus be conceived as the percentage reduction in the difference between the source and target cell states. While this may not correspond literally to the efficiency of a real-world experimental network perturbation, it is a very useful heuristic to estimate the relative effectiveness of different transcription factor perturbations. It is also instrumental in identifying an optimal number of transcription factors to perturb. As more transcription factors are predicted for perturbation, the precision score gradually increases up to a point at which it starts to level off with the inclusion of further perturbed transcription factors. This indicates the point at which further transcription factor perturbations may have less of an effect. It can also be a sign of low model confidence in the predicted perturbation as the model is unable to find another transcription factor to perturb that would have a strong effect. In cases where PrecICE does not predict any transcription factor with a precision score greater than 10-15%, it is likely because the model was unable to find a solution.

### Simplifying assumptions

The PreciCE algorithm uses many simplifying assumptions:

1. Transcription factors can regulate different genes in different cell types. While the PreciCE algorithm can technically be run on any underlying transcriptional network, we do not recommend using a “universal transcriptional network”. Instead, under the optimal settings, the PreciCE algorithm reconstructs its transcriptional networks solely from the input data of starting and target cells. This is both a strength (potentially greater network accuracy, no input network required) and a weakness (too few cells in input data translated into noisy networks). In cases where generation of such networks is difficult due to insufficient data or computational resources, we also provide pre-computed networks for commonly studied cell types.
2. The network edges represent transcription factor binding to proximal promoters. Enhancers are not included.
3. Edge weights between transcription factors and their target genes represent a linear relationship. In other words, the algorithm assumes that increasing the expression level of a transcription factor will linearly increase the expression level of its target genes, both in cases of individual gene perturbation effects as well as in combination This is not always the case in reality.
4. The algorithm is designed such that the effect of a transcription factor’s perturbation will propagate a defined number of steps (nodes) in the network, like ripples on a surface. This is not always true in reality.
5. The gene regulatory network inference is not perfect. For example, in our pluripotent-to-mesoderm prediction, *MESP1* has an out-degree of 0 (meaning *MESP1* is falsely predicted to regulate 0 genes). Still, our experimental data and that of others show that *MESP1* can promote mesoderm formation. In our pluripotent-to-mesoderm prediction, *MESP1* is still highly ranked because it is highly differentially expressed, and the model predicts that, after having modulated the higher-ranked genes, any further large network perturbations could be detrimental, so it prioritizes highly differentially expressed genes with low out-degree.

### Considerations for running the PreciCE algorithm

- Transformations of the input gene expression space beyond standard preprocessing, such as in the case of batch correction, should be avoided whenever possible because they risk impacting the true gene expression variation between the source and target cell states. However, in cases where experimental covariates vary between the source and target cell states, batch correction would be required regardless.
- Identification of cell state clusters is critical and care should be taken to avoid excessive heterogeneity in these clusters.
- When running the model, both differentially expressed and non-differentially expressed genes are included and add constraints to the model output.
- While some of the simplifying assumptions in PreciCE limit its flexibility to detect complex effects, they also enable it to be very confident in its prediction in case a clear solution exists. This would be reflected in high values of the precision score (>20%) while perturbing only a few transcription factors.

### Visualization of the PreciCE algorithm’s reconstructed network

The PreciCE algorithm’s reconstructed transcriptional networks were visualized as force-directed graphs using Observable (https://observablehq.com).

### Comparison of predictions for three different datasets

For comparisons of the PreciCE output from datasets Friedman et al., Pijuan-Sala et al., and Magnusson et al. (this study), we used a gene regulatory network inferred from the Friedman et al. dataset.

---

**Figure S1.**
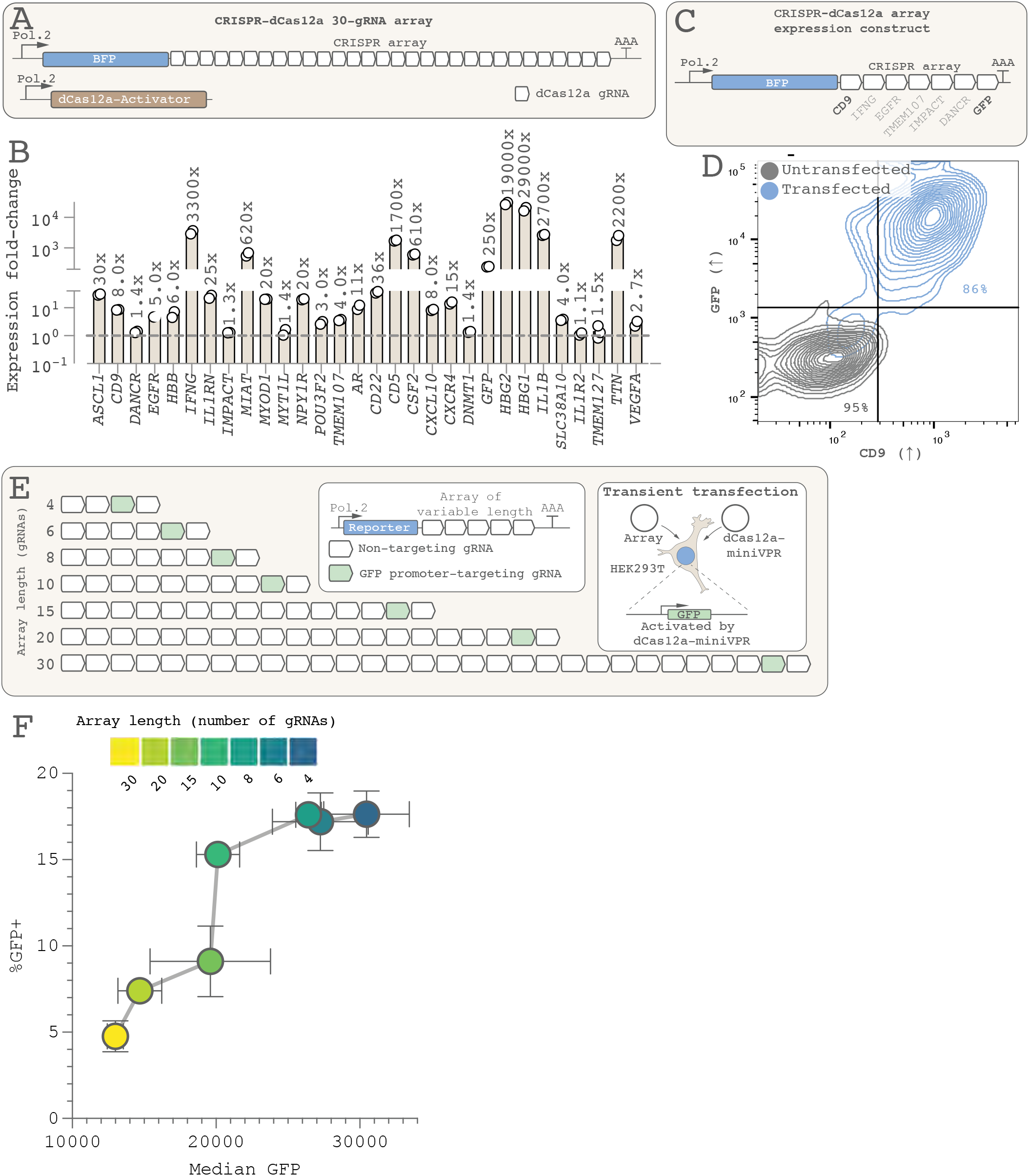
Engineered CRISPR-dCas12a arrays can be used for highly multiplexed gene control. **(A)** We expressed a 30-gRNA dCas12a array, preceded by a BFP reporter gene, under the control of a Pol. II promoter. Each gRNA targets the promoter of one gene for upregulation. (**B**) When this construct and a dCas12a-miniVPR activator are co-transfected in HEK293T cells carrying genomically integrated GFP, all 30 target genes are upregulated, as measured by RT-qPCR. Expression fold-change varies widely and depends in part on the efficiency of the gRNA and the baseline expression of the target gene. (**C**) To estimate the efficiency of multi-gene modulation on the single-cell level, we expressed a 7-gRNA CRISPR array and analyzed two target gene products (CD9 and GFP) by flow cytometry 48 hr later. (**D**) Co-activation of these two genes is highly efficient, indicating that multi-gene activation occurs on the single-cell level. (**E**) To evaluate whether there is an efficiency drop as CRISPR arrays grow longer, we generated seven CRISPR arrays of varying length (4-30 gRNAs) where one gRNA (green) targets the GFP promoter for activation and all other gRNAs (white) contain non-targeting spacers, transfecting this construct together with a dCas12a-miniVPR activator. (**F**) Increasing the number of gRNAs in the array reduces GFP activation, possibly due to dilution of available dCas12a protein (error bars show standard deviation). Note that not only does the median GFP fluorescence level decrease but also the percentage of GFP-expressing cells (despite gating for cells that had taken up both constructs). However, even with a 30-gRNA array, there is only a ∼50% drop in median GFP activation efficiency and a ∼70% drop in the percentage of GFP^+^ cells, supporting the use of Pol. II-transcribed CRISPR arrays for highly multiplexed gene regulation in human cells. We deliberately used a weaker dCas12a construct (wild-type protein sequence rather than the improved hyper-dCas12a) to increase sensitivity and dynamic range.

**Figure S2.**
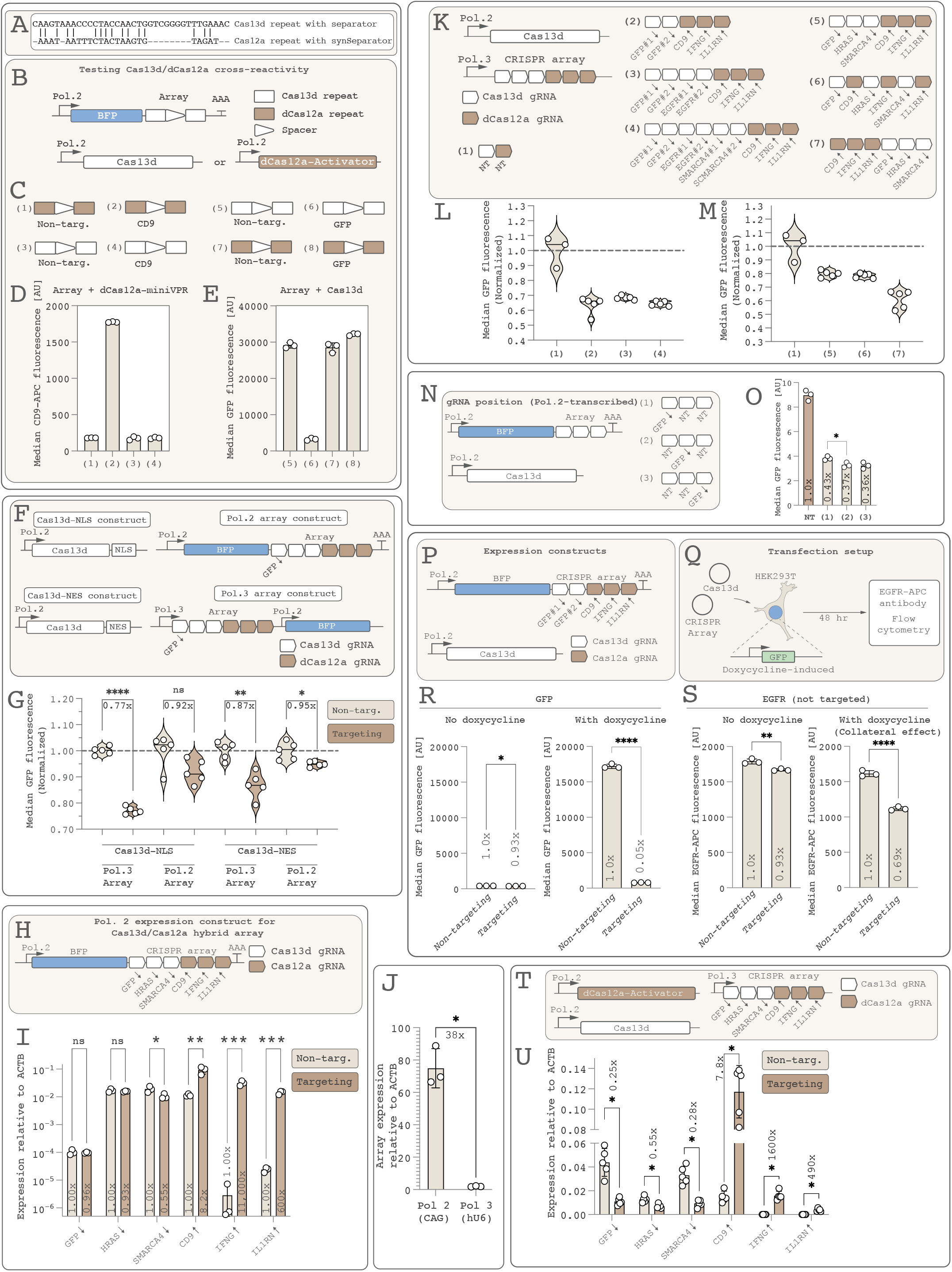
Optimization of the CRISPR-Cas13d/dCas12a hybrid array system. **(A)** The gRNA repeats of Cas13d and Lb-dCas12a, including separator sequences^20^, are very different from one another. (**B**) To test Cas13d/dCas12a cross-reactivity, we expressed a CRISPR “array” consisting of a single gRNA with either a Cas13d repeat or a dCas12a repeat, a spacer (targeting CD9 for upregulation or GFP for downregulation), and another repeat, all preceded by a BFP gene and expressed under a Pol. II promoter (CAG). This array construct was co-transfected with either Cas13d or dCas12a-miniVPR. (**C**) We designed eight such CRISPR arrays combining either the Cas13d or dCas12a repeat with spacers targeting the CD9 promoter ((2), (4)) or GFP gene body ((6), (8)) or corresponding non-targeting spacers ((1), (3), (5), (7)) (Diagrams show only the array portion of the constructs). (**D**) We transfected arrays (1)-(4) together with a dCas12a-miniVPR activator in HEK293T cells, stained cells with a CD9-A467 antibody 48 hr later and analyzed by flow cytometry. Only cells co-expressing the CD9-targeting gRNA containing a dCas12a repeat activated CD9, showing that dCas12a cannot use gRNAs containing Cas13d’s repeat. (**E**) Conversely, we used HEK293T cells that constitutively express genomically encoded GFP and transfected these with arrays (5)-(8) together with Cas13d. Only cells expressing the GFP- targeting gRNA containing a Cas13d repeat repressed GFP, showing that Cas13d does not recognize the repeat region of dCas12a’s gRNA. (**F**) We tested the effects of fusing Cas13d to a nuclear localization signal or nuclear export signal, and of expressing a hybrid Cas13d/dCas12a array under a Pol. II or Pol. III promoter to analyze GFP repression efficiency in HEK293T cells constitutively expressing GFP. (**G**) Cas13d-NLS together with a Pol. III-transcribed array works best, as measured by flow cytometry. (Note that the y axis starts at 0.70. Samples were normalized to their own non-targeting values. Each data point represents the median fluorescence of one experimental replicate, as measured by flow cytometry) (**H**) Accordingly, using a Pol. II expression construct to drive Cas13d/dCas12a hybrid arrays does not work well. (**I**) When co-transfected in HEK293T cells with Cas13d and dCas12a-miniVPR, gene repression is barely detectable, though dCas12a-miniVPR-mediated activation works well. (**J**) This is not due to low expression of the CRISPR array. In fact, Pol. II-mediated expression leads to much higher levels of the array transcript (using primers specific to the array) than expressing the array from a Pol. III promoter (see panel **F**). (**K**) To test the effects of gRNA position, CRISPR array length, and the number of unique gRNAs targeting a gene when the array is expressed under a Pol. III promoter, we designed multiple CRISPR array constructs and co-transfected each of these with Cas13d in HEK293T cells constitutively expressing GFP, measuring GFP fluorescence by flow cytometry 48 hr later but not analyzing other target genes. (**L**) CRISPR array length does not cause a measurable reduction in GFP repression efficiency, though we cannot exclude that this is because the GFP-targeting gRNAs are the first gRNAs in the array. (**M**) GFP repression works better when Cas13d gRNAs are encoded after dCas12a gRNAs on the hybrid array. Surprisingly, GFP repression still works when the array has an alternating Cas13d/dCas12a gRNA arrangement ((6)), despite Cas13d being unable to process dCas12a’s gRNAs (**E**). This suggests that Cas13d is functional even with a full 44-nt dCas12a gRNA protruding from its own gRNA’s 3’ end. GFP repression is more effective when using two unique gRNAs instead of just one (compare (2)-(4) with (5)). (Note that the same non-targeting array data are plotted in panels **L** and **M**). (**N**) We also tested positional effects in a Pol. II-transcribed array. (**O**) Any such effects were very subtle, consistent with previous results for Pol. II-transcribed Cas12a arrays^20^. Therefore, positional effects may be different depending on whether arrays are transcribed under a Pol. II or Pol. III promoter. (**P)** To test Cas13d collateral activity, we used a hybrid CRISPR-Cas13d/dCas12a array targeting (among other genes) GFP for repression. (**Q**) We co-transfected this with a Cas13d construct into HEK293T cells expressing doxycycline-inducible GFP and measured the expression level of the non-targeted gene product EGFR using flow cytometry (see Methods). (**R**) Doxycycline administration activates strong GFP expression and efficient Cas13-mediated GFP repression. (**S**) But it also leads to marked downregulation of the collateral marker gene EGFR, specifically in cells expressing the GFP-targeting gRNA. This effect, which we interpret as collateral activity, is much reduced when doxycycline was not administered and GFP expression is therefore much lower (S; “No doxycycline”), showing that collateral activity depends on expression level of the target gene. (**T**) These tests informed the design of our final CRISPR- Cas13d/dCas12a hybrid array, which we tested in HEK293T cells carrying genomically integrated, doxycycline-inducible GFP (but relying on leaky GFP expression in the absence of doxycycline administration to avoid collateral activity). (**U**) Three genes can be repressed, and three genes simultaneously activated, as measured by RT-qPCR 48 hr after transfection. All error bars represent standard deviation.

**Figure S3.**
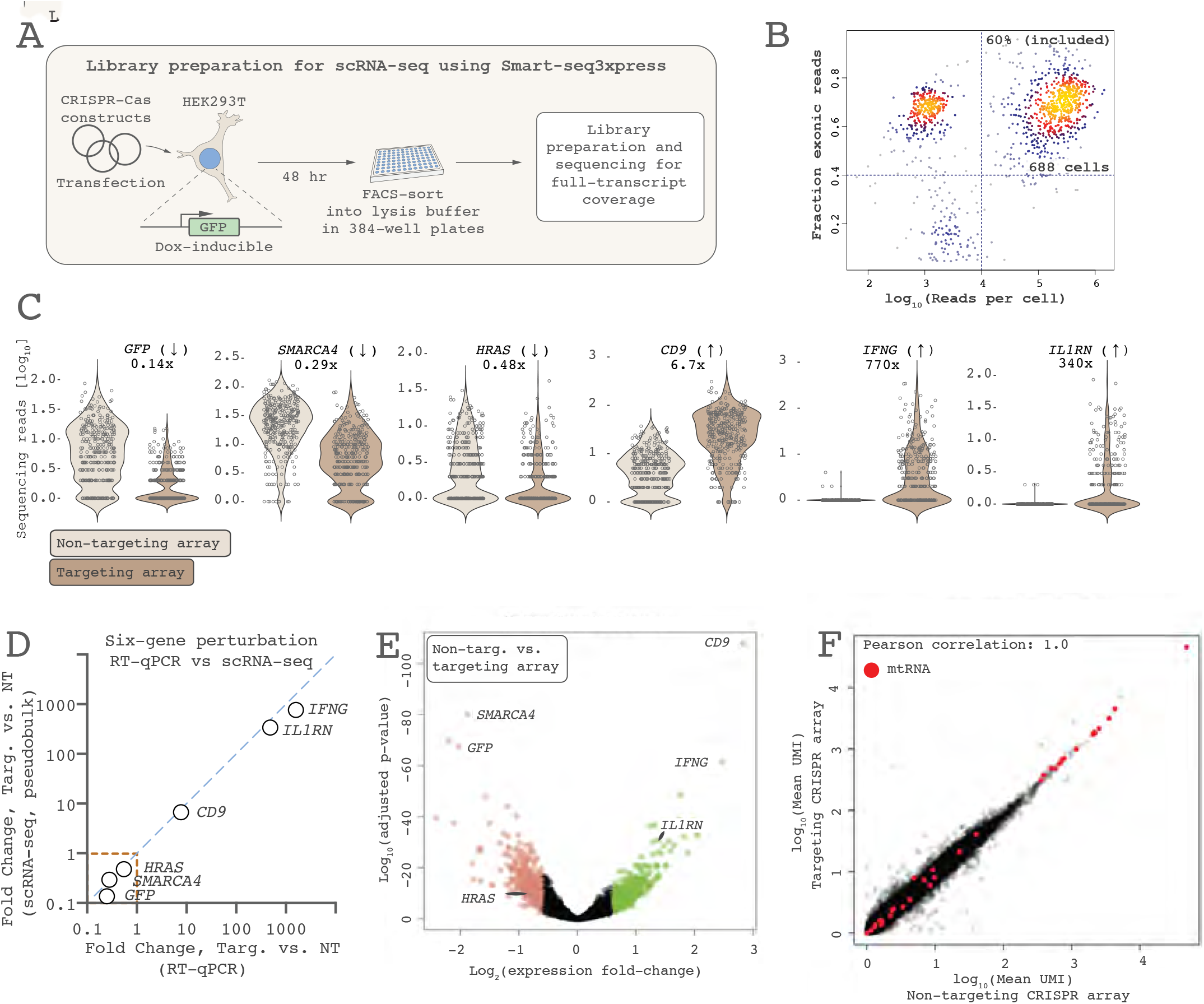
ScRNA-seq analysis of multi-gene perturbation using the CRISPR- Cas13d/dCas12a hybrid array system. **(A)** We transfected HEK293T cells with a dCas12a- miniVPR, Cas13d, and a Pol. III-encoded Cas13d/dCas12a hybrid array (see Fig. 2A). We relied on leaky GFP expression without doxycycline administration to prevent Cas13d collateral activity caused by high target gene expression. Cells were FACS-sorted into 384-well plates and processed for sequencing using the Smart-seq3xpress workflow^26^. (**B**) Post-sequencing quality control plot of scRNA-seq libraries, showing that the final dataset consisted of 688 cells. (**C**) Violin plots demonstrating simultaneous repression of *GFP*, *SMARCA4*, and *HRAS*, and activation of *CD9*, *IFNG*, and *IL1RN* (Each data point is one cell. Expression fold-change values correspond to the mean read count across cells expressing the targeting array divided by that of cells expressing the non-targeting array.) (**D**) Expression fold-change values correspond well between RT-qPCR and scRNA-seq (dashed line delineates Cas13d target genes). (**E)** The six target genes are among the most highly differentially expressed genes in these cells, though many other genes are differentially expressed at this 48 hr time point, possibly because of broad secondary effects for some of the target genes. (**F**) Differential expression of mitochondrial transcripts can be used to estimate Cas13d collateral effects, as these transcripts are protected from Cas13d by the mitochondrial membrane and will artifactually appear upregulated when collateral effects are present^25^. However, cells expressing the CRISPR array with targeting gRNAs show no apparent upregulation of mtRNAs (red data points), showing that collateral cleavage is not detected.

**Figure S4.**
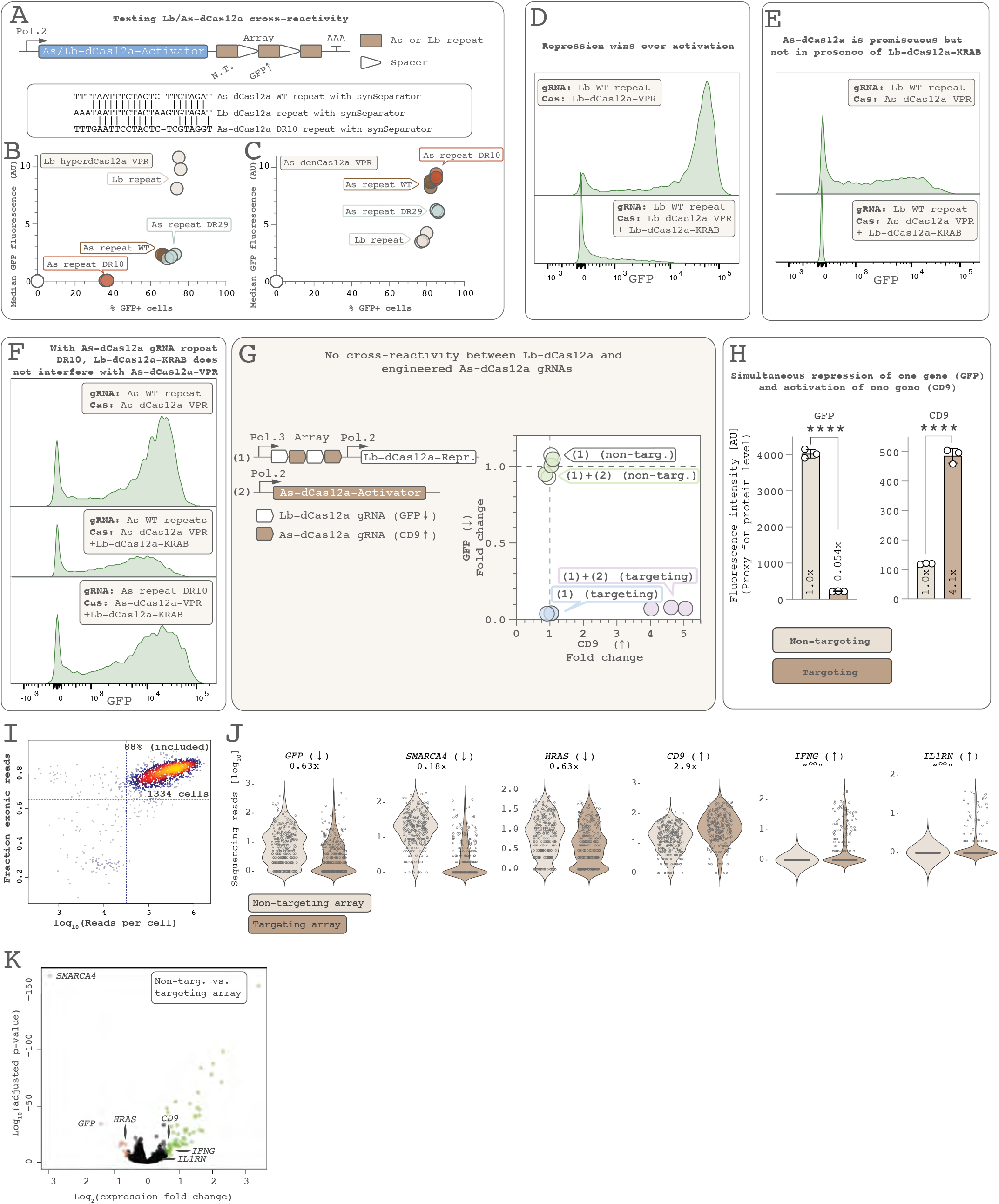
Development of a CRISPR Lb-dCas12a/As-dCas12a hybrid system. (**A**) To test cross-reactivity between Lb-dCas12a and As-dCas12a, we transfected HEK293T cells with one plasmid expressing either As-dCas12a-VPR or Lb-dCas12a-VPR followed by a CRISPR array. The array contained two gRNAs, one with a non-targeting spacer and the other targeting the promoter (TRE3G) of GFP (genomically integrated in the HEK293T cells) for GFP activation. The array’s gRNAs contained either Lb-dCas12a repeats or wild-type As-dCas12a repeats, allowing us to test combinations of dCas12a and gRNA repeat variants. We also tested two engineered variants (DR10, DR29) of the As-dCas12a gRNA repeats^32^. Sequence alignment shows how the DR10 variant is less similar than the wild-type As-dCas12a repeat to the Lb-dCas12a repeat (also shown are AAAT/TTTT/TTTG synSeparators^20^. These constructs were transfected in HEK293T cells containing genomically integrated TRE3G-GFP; GFP fluorescence was measured by flow cytometry 48 hr later. Median GFP fluorescence was plotted against the percentage of GFP+ cells for a more nuanced view of GFP activation strength than either metric alone. (**B**) Lb-dCas12a- VPR works best with its own gRNA repeat. Lb-dCas12a shows some undesired promiscuous activity also with gRNAs containing the As-dCas12a WT and DR29 repeats. But, promisingly, Lb- dCas12a shows very little cross-reactivity with the As-dCas12a DR10 repeat. (**C**) As-dCas12a- VPR is slightly more promiscuous than Lb-dCas12a-VPR and can activate GFP using a gRNA containing Lb-dCas12a’s repeat. Fortuitously, the DR10 repeat performs at least as well as the wild-type As-dCas12a repeat. Thus, the DR10 variant is a crucial component for enabling non-interference between Lb- and As-dCas12a. (**D**) To enable further optimization, we tested what happens when a dCas12a-Activator and dCas12a-Repressor are recruited to the same site on a target gene promoter. First, an Lb-dCas12a-VPR-Activator, when expressed alone together with a GFP-targeting Lb-dCas12a gRNA, activates genomically integrated GFP (top histogram), as expected. But if co-expressed with the same dCas12 variant (Lb-dCas12a) fused to a repressor (KRAB), GFP activation no longer occurs (bottom histogram). This shows that when an activator and a repressor are recruited to the same site, repression wins. (**E**) This holds true also in competition between As-dCas12a and Lb-dCas12a: An As-dCas12a-VPR activator promiscuously uses a gRNA containing an Lb-dCas12a repeat to activate GFP (top histogram). But when As-dCas12a-VPR is co-expressed with an Lb-dCas12a-KRAB repressor, As-dCas12a- VPR is no longer able to promiscuously activate GFP. Panels **D-E** thus show that the repressor domain should be on the less promiscuous of the two dCas12a variants (i.e., Lb-dCas12a) to prevent undesired promiscuous repression of target genes intended for activation. (**F**) When As- dCas12a uses the DR10 repeat for its gRNAs, Lb-dCas12a is by far the less promiscuous of the two dCas12a variants. Indeed, even if Lb-dCas12a-KRAB promiscuously interferes with As- dCas12a-VPR when the wild-type As-dCas12a repeat is used, and Lb-dCas12a-KRAB thus prevents As-dCas12a-VPR from fully activating GFP (top two histograms), such undesired interference is no longer seen when the As-dCas12a gRNAs use the DR10 repeat variant (bottom histogram). Thus, to prevent undesired interference between Lb- and As-dCas12a, the following design should be used: As-dCas12a gRNAs should use the DR10 repeat variant; Lb-dCas12a should carry the repressor domain while As-dCas12a carries the activator domain; all gRNAs should be expressed together with the Lb-dCas12a-Repressor gene and not with the promiscuous As-dCas12a gene. (**G**) When this is true, co-expression of the two dCas12a constructs (“(1)” and “(2)”) enables simultaneous activation of an As-dCas12a-Activator target gene (CD9) and repression of an Lb-dCas12a-Repressor target gene (GFP) in a HEK293T cell line that constitutively expresses GFP (different cell line from the one in panels **A-F**). In cells that only take up the Lb-dCas12a-Repressor + CRISPR array construct (“(1)”), repression of the Lb-dCas12a target gene GFP occurs but no promiscuous repression of the As-dCas12a target gene CD9, demonstrating orthogonality (The plot shows median fluorescence levels in experimental triplicates, each sample normalized to the average of its non-targeting control). (**H**) shows experimental replicates of the FACS plot shown in Fig. 2E. (**I**) Quality control plot of scRNA-seq (Smart-seq3xpress) data for HEK293T cells expressing the CRISPR array in Fig. 2F. (**J**) Subsetting for cells that express both dCas12a variants, violin plots show the simultaneous downregulation of *GFP*, *SMARCA4*, and *HRAS*, and upregulation of *CD9*, *IFNG*, and *IL1RN* (Each data point is one cell. Fold-change corresponds to the mean read count across cells expressing the targeting array divided by that of cells expressing the non-targeting array. Fold-change could not be calculated for *IFNG* and *IL1RN* as they are not normally expressed in these cells.) (**K**) A volcano plot of scRNA-seq data shows that the Lb-dCas12a-Repressor target genes are among the most highly differentially expressed genes, but the As-dCas12a-Activator target genes are more subtly modulated. At this 48-hr time point, a number of secondary transcriptional effects have taken place. (Note that the limma/voom algorithm computes expression fold-change differently from panel **J**). All error bars represent standard deviation.

**Figure S5.**
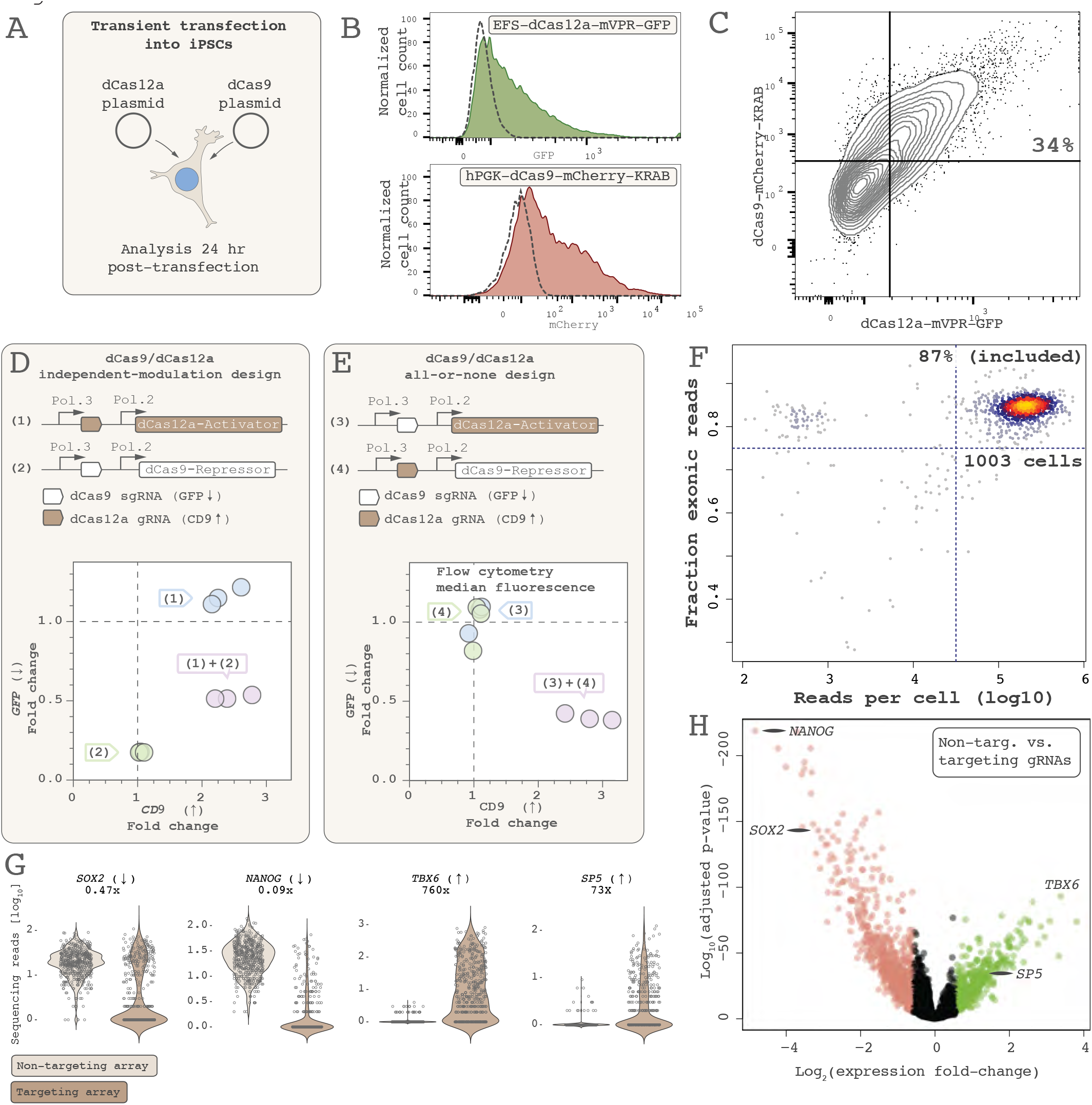
A dCas9/dCas12a two-construct architecture for all-or-none gene modulation. (**A**) In transfection experiments, some cells may fail to take up both of two co-transfected plasmids. We co-transfected iPSCs with two plasmids expressing dCas12a-miniVPR-GFP and dCas9-mCherry-KRAB, respectively. (**B-C**) Twenty-four hours later, a third of the cells express both constructs. (**D**) If a dCas12a-Activator and dCas9-Repressor gene are encoded on the same plasmid as their own gRNAs (“(1)” and “(2)”), cells transfected with only one construct will perform only gene activation [CD9*↑*] or only gene repression [*GFP↓*] (in HEK293T cells constitutively expressing genomically integrated GFP), which risks introducing undesired heterogeneity in cell fate programming applications. Cells that take up both constructs simultaneously activate CD9 and repress GFP. (Each data point represents median fluorescence in one flow cytometry replicate, each sample normalized to the average of its non-targeting control). (**E**) But by encoding gRNAs from each dCas protein onto the expression vector of the other dCas protein, only cells that take up both constructs (“(3)+(4)”) simultaneously upregulate CD9 and downregulate GFP. Cells that take up only one of the two constructs (“(3)” or “(4)”) do nothing. This architecture constitutes a logical AND gate, ensuring that only cells that take up construct (3) AND construct (4) perform any gene modulation. (**F**) Post-sequencing quality control plot of scRNA-seq libraries of FACS-sorted iPSCs expressing a dCas12a-miniVPR activator and dCas9-KRAB repressor construct encoded with the gRNAs shown in Fig. 2I, showing that the final dataset consisted of 1003 cells. (**G**) Violin plots of the data shown in Fig. 2I (Each data point represents one cell. Fold-change corresponds to the mean read count across cells expressing the targeting gRNAs divided by that of cells expressing the non-targeting gRNAs). (**H**) Volcano plot showing that the four target genes are among the most highly differentially expressed genes, though massive global transcriptional changes have also occurred 48 hr after perturbing these four transcription factors.

**Figure S6.**
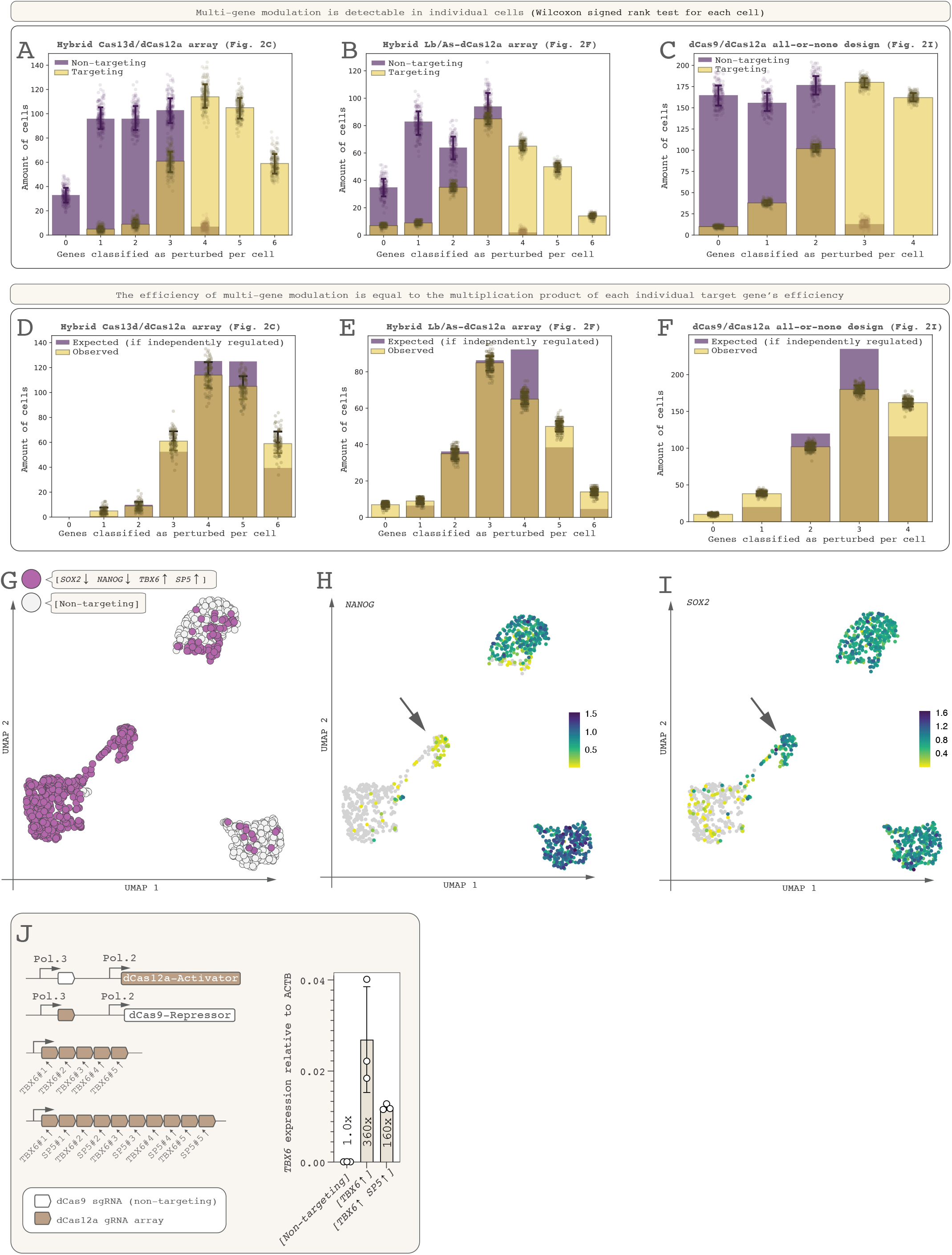
Estimating multi-gene regulation in individual cells. With CRISPR-mediated gene perturbation, target genes do not show binary on/off modulation but gradual differences in expression levels (compare violin plots in Figs. S3C, S4J, S5G). Moreover, scRNA-seq data is noisy due to transcriptional bursting and dropout effects. (**A-C**) We sought to assess whether multi-gene regulation occurs in individual cells. We used the following method (see **Methods**). For each target gene, we used the control cells (expressing the non-targeting gRNAs) as reference for the baseline expression pattern of that gene. Based on that, we set an expression threshold, beyond which that gene was considered “perturbed”. That threshold was arbitrarily set to include as many as possible of the cells expressing the targeting gRNAs and as few as possible of the control cells expressing the non-targeting gRNAs. Then, for every individual cell, we asked how many of the target genes were thus “perturbed” and plotted the results in panels A-C (data points represent bootstrapping iterations and standard deviation). The results are likely an underestimate because transcriptional bursting and dropout effects introduce much noise and make the data zero-inflated. None of the target genes entirely show an “off/on” pattern (e.g., zero in all control cells; high in all perturbed cells). An indication that it is difficult to set strict perturbation thresholds can be seen when looking at the data for the control cells (expressing the non-targeting array): for many of these cells, some target genes were statistically classified as “perturbed”. Such erroneous classification was likely caused, at least in part, by such noise. (**D-F**) Despite such noise, our data allowed us to analyze whether there were intrinsic biases for or against multi-gene regulation inside each cell. For each of the target genes, some fraction of cells will be classified as “perturbed” given the perturbation thresholds described above. If the target genes are independently regulated, the number of cells showing all genes perturbed is equal to the multiplication product of all genes’ individual perturbation fractions. For example, in a hypothetical scenario where there are six target genes and, for each gene, 50% of cells are classified as “perturbed”, then the fraction of cells showing all six genes perturbed will be 0.5^6^=0.016 if the six genes are independently modulated. Corresponding fractions can be calculated to estimate how many cells should show 5, 4, 3, 2, 1, and 0 genes perturbed. If, on the other hand, cells expressing a CRISPR array show any biases toward regulating all target genes or only subsets of target genes, the experimentally observed data should deviate from that independent-regulation scenario. We used the perturbation thresholds defined for each target gene and performed this calculation. Indeed, the experimentally observed data matched the independent-regulation scenario remarkably well (data points show 100 bootstrapping iterations and error bars represent standard deviation). The data suggest there is a slight desirable bias toward perturbing all target genes. Importantly, there is no bias *against* perturbing multiple genes per cell. These data suggest that the efficiency of multi-gene perturbation on the single-cell level is a direct product of the perturbation efficiencies of each individual target gene. (**G**) With [*TBX6*↑ *SP5*↑ *SOX2↓ NANOG↓*] perturbation using the dCas9/dCas12a design (Fig. 2G**-I**), cells have separated into different clusters in a UMAP 48 hr after transfection. (**H**) *NANOG* downregulation is more effective than *SOX2* downregulation (**I**; compare **Fig. S5G**). Because perturbation affects not only perturbation magnitude per cell but also the fraction of cells that execute the perturbation (see **Fig. S1F**), a subset of cells shows *NANOG* downregulation but not *SOX2* downregulation (**H-I**, arrows). (**J**) Perturbation efficiency is affected in part by the number of CRISPR target genes: *TBX6* upregulation is more efficient when only *TBX6* is targeted than when both *TBX6* and *SP5* are targeted. The difference in perturbation magnitude is consistent with dilution of available dCas12a protein but can be rescued by boosting dCas12a expression level (see **Fig. S12A-B**).

**Fig. S7.**
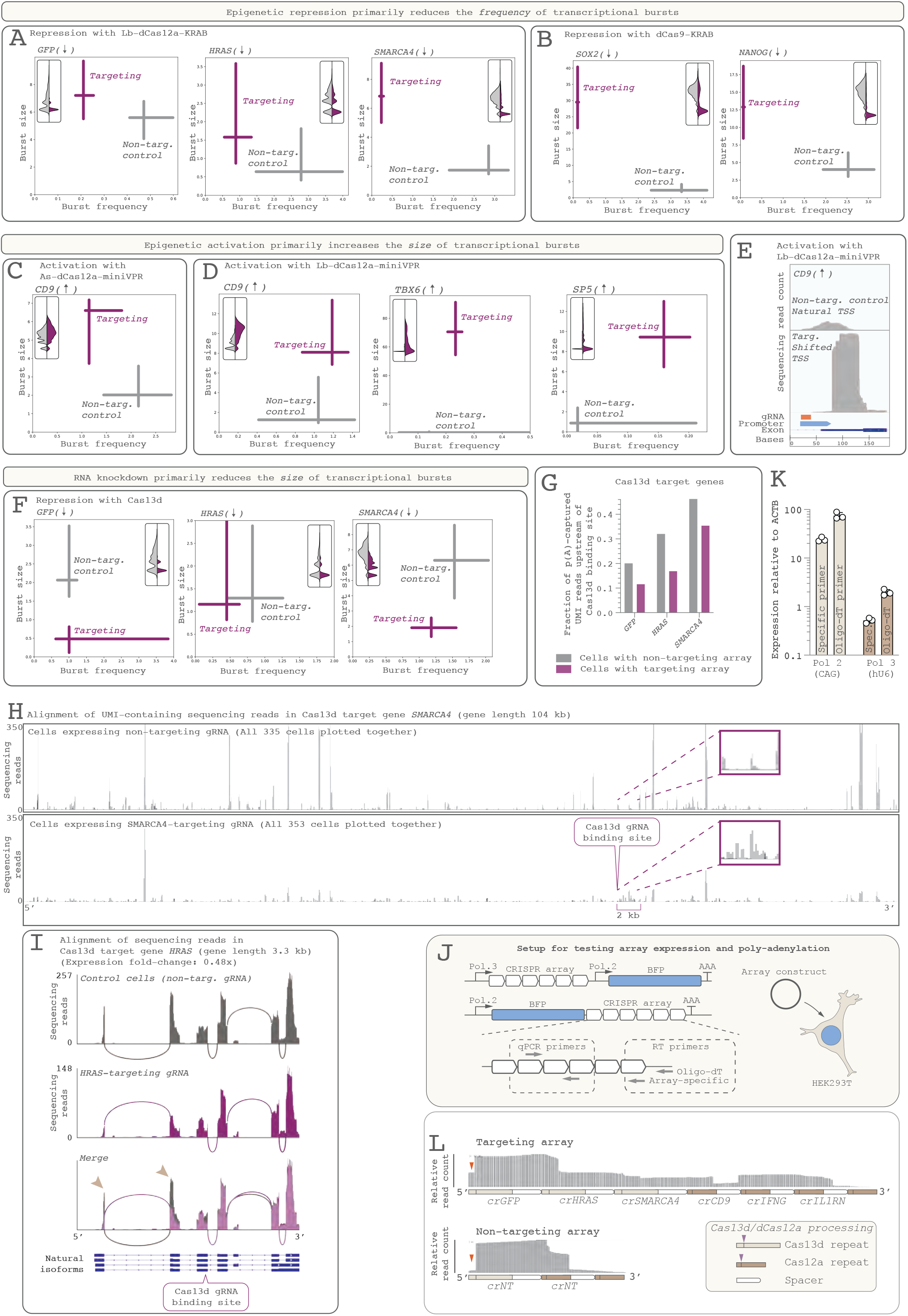
Mechanisms of CRISPR-based gene activation and repression. (**A-B**) CRISPRi with dCas12a-KRAB (**A**) and dCas9-KRAB (**B**) achieves its repressive effect through reduction of burst frequency. The data suggest a simultaneous *increase* in burst size, perhaps as a compensatory mechanism. Insets show violin plots of UMI-containing read count for cells expressing non-targeting (gray) and targeting (purple) gRNAs for comparison. Error bars represent 95% confidence interval. (**C-D**) Gene upregulation with dCas12a-miniVPR happens mainly through increased burst size (Target genes *IFNG* and *IL1RN* could not be analyzed because baseline expression was 0). (**E**) For the CRISPRa target gene *CD9*, the dCas12a gRNA is located within the natural transcriptional start site (plots represent alignment of UMI-containing sequencing reads, zoomed in to the start of the first exon). Likely because of steric hindrance, CRISPRa-induced transcription starts ∼80 bp downstream of the normal start site (“Promoter”: UCSC Genome Browser EPDnew track). (**F**) Cas13d-targeted transcripts show reduced burst size but not frequency, consistent with Cas13d’s role as an mRNA-targeting enzyme. Note that *HRAS* repression was so inefficient that the bursting inference yielded non-significant differences. (**G**) We investigated whether Cas13d-targeted partial transcripts linger long enough to be detectable. Because Smart-seq3xpress uses an oligo-dT primer to capture poly-adenylated RNAs during reverse transcription, Cas13d-targeted transcripts might show fewer sequencing reads aligning upstream (5’) of the Cas13d binding site if Cas13d degradation products linger in the cell after Cas13d-mediated transcript cutting. We first analyzed all control cells expressing the non-targeting CRISPR array to see what fraction of UMI-containing sequencing reads normally align upstream of the site where targeting Cas13d gRNAs would bind (gray bars). Next, analyzing cells expressing the targeting gRNAs (purple bars), we found that a lower fraction of UMI-containing sequencing reads align upstream of the Cas13d binding site than in the control cells (gray bars). This suggests that at least the 3’ end of the Cas13d-targeted transcript lingers long enough to be detectable (This method cannot be used to detect any putative lingering 5’ end). (**H**) In support of this, the long Cas13d target transcript *SMARCA4* (105 kb) has a single region where cells expressing the *SMARCA4*-targeting gRNA show more UMI-containing sequencing reads than control cells expressing the non-targeting gRNA, namely the 2-kb region immediately downstream of the Cas13d binding site (inset). This suggests that the 3’ part of the Cas13d-degraded transcript did indeed linger in the cell such that the UMI was added there during reverse transcription. Furthermore, it suggests that Cas13d can cut a transcript indiscriminately somewhere within 2 kb of its target site. This pattern was not visible in the target transcripts *GFP* (∼1 kb) or *HRAS* (∼3 kb), possibly because it takes a longer transcript to make this pattern visually apparent. (**I**) Sashimi plot of the downregulated Cas13d target gene *HRAS* (expression fold-change 0.48x), where lines show exon-spanning sequencing reads. Note that, in addition to an overall reduction in *HRAS* transcript abundance (note difference in scale for y-axes), many transcripts appear to lack the first two exons (arrowheads), suggesting that Cas13d could be used to generate synthetic isoforms (though these likely lack a 5’ UTR). (**J**) Experimental setup to address whether Pol. III-transcribed transcripts get poly-adenylated. For reverse transcription, we used primers that were either specific to one gRNA sequence or oligo-dT primers that would only amplify the array if it was poly-adenylated. We used RT-qPCR primers specific to a region of the CRISPR array upstream of the RT primers. A Pol. II-transcribed array construct was used as a positive control, as we know this gets poly-adenylated. The array constructs (without Cas genes) were transfected into HEK293T cells as two separate experimental conditions. (**K**) Both array-specific and oligo-dT RT primers amplify the Pol. II-transcribed array, as expected, but surprisingly also the Pol. III-transcribed array, indicating that this has become poly-adenylated. (**L**) Accordingly, Smart- seq3xpress sequencing reads (from the Cas13d/dCas12a CRISPR array experiment, **Fig. 2C**) align to the CRISPR arrays. Inset shows where on the gRNA repeats the respective Cas proteins cut the gRNAs to process the CRISPR array. Note that the aligned sequencing reads indicate that CRISPR array processing has taken place (orange arrowheads) but that processing has not proceeded to completion, suggesting that array transcripts were in excess. Note that Cas13d array processing is more efficient for the shorter non-targeting array (1 Cas13d gRNA) than the longer targeting array (3 Cas13d gRNAs), also suggesting Cas13d protein availability may make array processing a limiting step for long CRISPR arrays. All error bars represent standard deviation.

**Figure S8.**
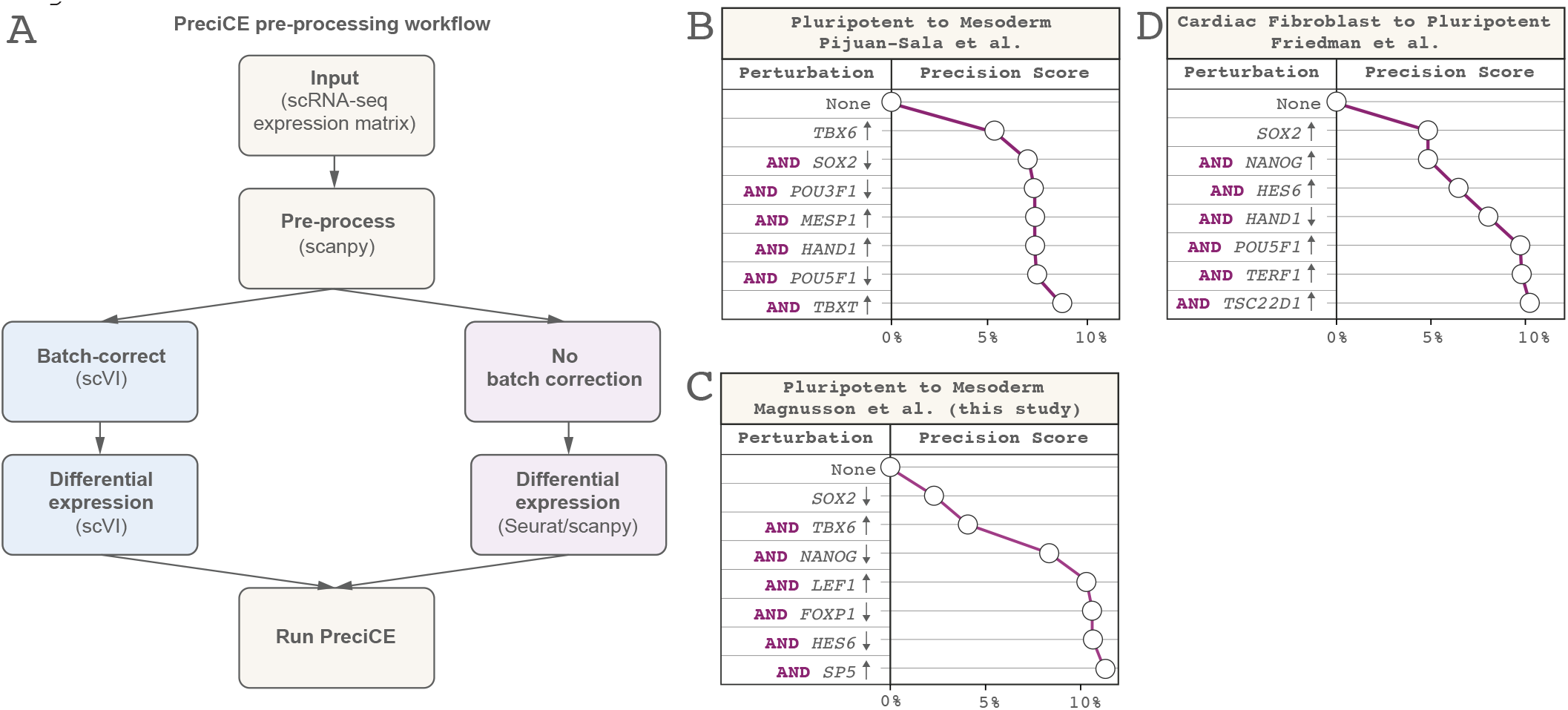
Supporting analyses for the PreciCE algorithm. **(A)** Diagram illustrating the PreciCE workflow: Single-cell RNA sequencing data is first preprocessed using Scanpy. If the source and target cell states come from different experimental batches, users can opt to run batch effect correction with scVI. Next, differential expression analysis is performed between the source and target cell states, using scVI for batch-corrected data, or Seurat/Scanpy for uncorrected samples. The resulting data is then fed into the PreciCE model to rank transcription factor perturbations. (**B-C)** Predictions for the conversion of pluripotent stem cells to mesoderm cells using scRNA-seq data of mouse in vivo embryonic differentiation **(B)** and human in vitro iPSC-to-mesoderm differentiation using a small-molecule protocol (this study; **C**), using the transcriptional network reconstructed from the Friedman et al. dataset. **(D)** To our knowledge, no datasets exist for validating simultaneous up- and downregulation, but for the reprogramming of fibroblasts to pluripotent stem cells, the PreciCE algorithm predicts multiple genes genes known to be important for pluripotent stem cells (*SOX2, NANOG, POU5F1/OCT4*).

**Figure S9.**
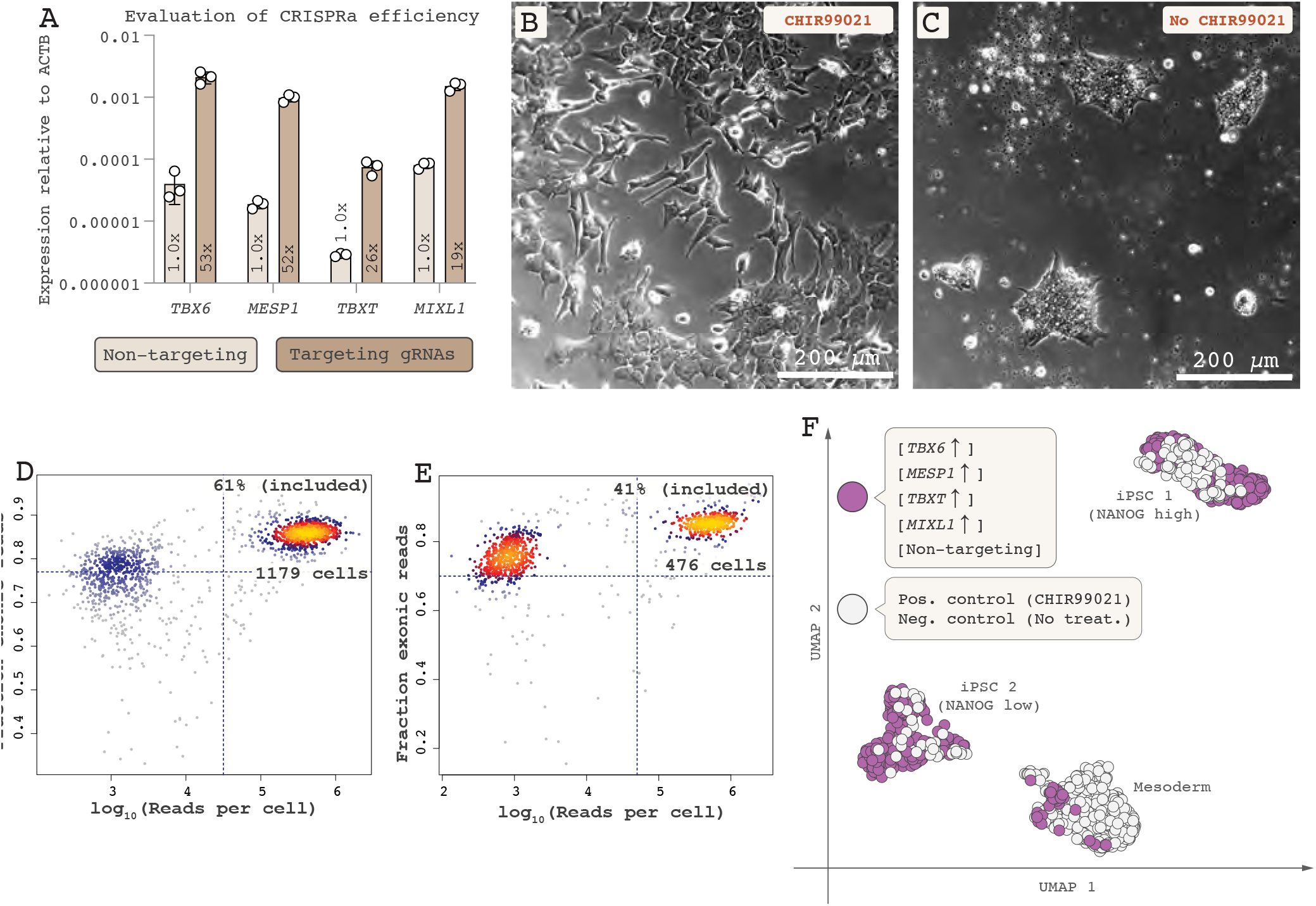
Experimental testing of the PreciCE algorithm’s top-ranked gene *TBX6* for iPSC-to-mesoderm conversion. (**A**) Single-gene CRISPRa of *TBX6*, *MESP1*, *TBXT*, or *MIXL1* shows activation by 19-53-fold in iPSCs 48 hr post-transfection (Error bars represent standard deviation). (**B**) The positive-control protocol using the Wnt-activating small molecule CHIR99021 differentiates iPSCs to mesoderm cells in two days, with a morphology that would later be seen in cells differentiated using our CRISPR-based systems. (**C**) Colonies of undifferentiated iPSCs. (**D-E**) Post-sequencing quality control of FACS-sorted cells from single-gene CRISPRa for mesoderm conversion (**D**) and from the small-molecule positive control condition (**E)**. Successful sequencing libraries are found in the top right quadrant. Note that the fraction of high-quality libraries is low (61% and 41%, respectively) due to unexpectedly inefficient FACS sorting into 384-well plates. (**F**) The resulting scRNA-seq data was plotted in a UMAP together with cells from a small-molecule-based (CHIR99021) mesoderm differentiation protocol as positive control.

**Figure S10.**
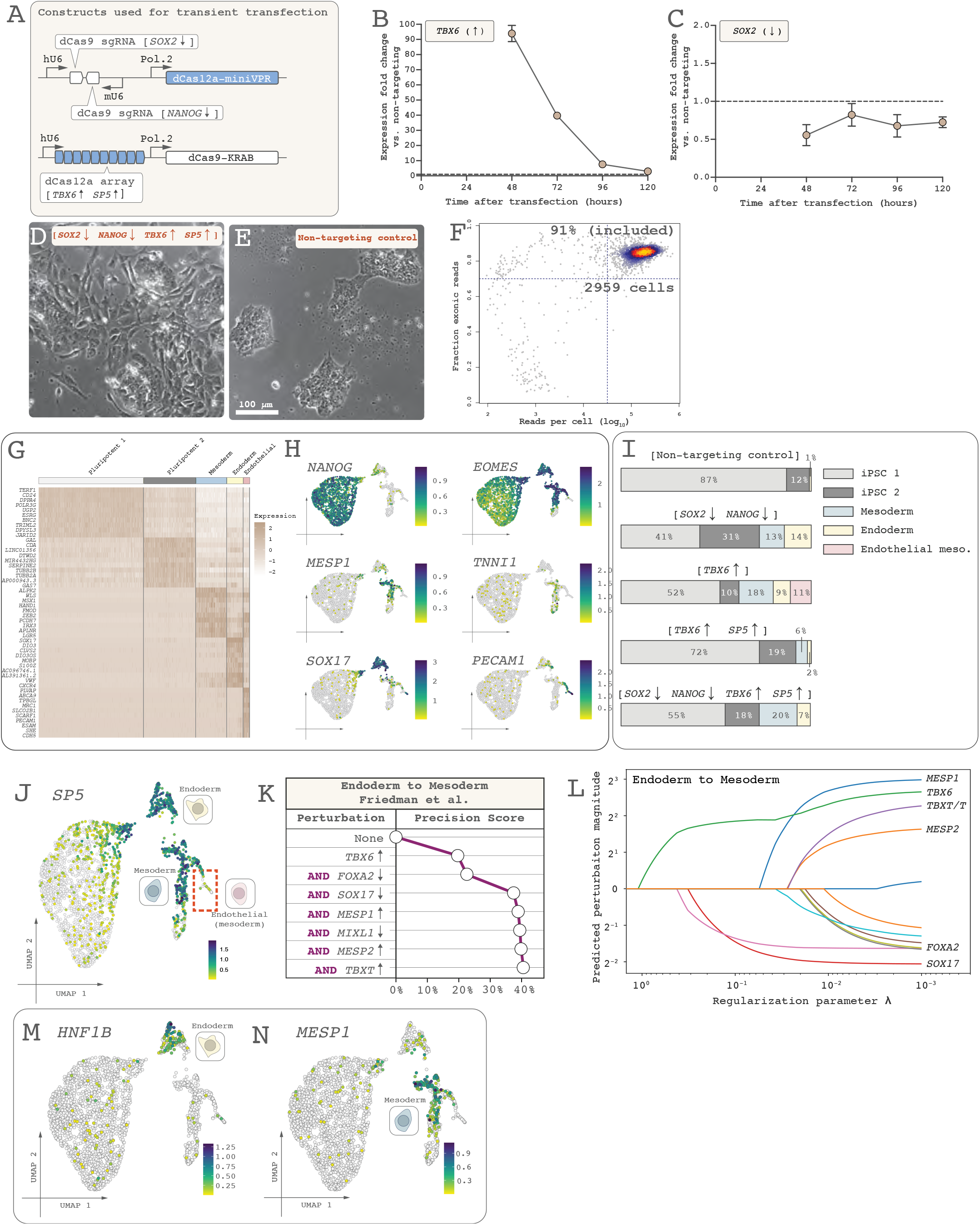
Cell fate logic using PreciCE for mesoderm differentiation. **(A)** We used transient transfection of our dCas9/dCas12a all-or-none design for these experiments, as this system allows rapid design-build-test cycles with minimal cell-to-cell heterogeneity. (**B**) Transient transfection of plasmids encoding the dCas9/dCas12a system (**Fig. 2G**) leads to miniVPR- mediated target gene activation (*TBX6*) that peaks 48 hr after transfection and then drastically declines (Graph represents RT-qPCR data of HEK293T cells; error bars represent standard deviation of three replicates). (**C**) dCas9-KRAB-mediated target gene repression (*SOX2*) is longer-lasting, lasting at least 5 days. (**D-E**) Light microscopy images of cells 96 hr post-transfection show morphological changes consistent with differentiation (**D**) compared to cells transfected with non-targeting gRNA (**E**). (**F**) A post-sequencing quality control plot shows that 91% of sequencing libraries (2959 cells) were of high quality and were used for downstream analysis. (**G**) Global data structure of iPSCs subjected to five different perturbations ([*TBX6*↑], [*TBX6*↑ *SP5*↑], [*SOX2↓ NANOG↓*], [*TBX6*↑ *SP5*↑ *SOX2↓ NANOG↓*], [Non-targeting]), showing top genes defining each cluster from **Fig. 5A**. (**H**) Marker genes showing the presence of iPSCs (*NANOG*), early differentiating iPSCs and mesendoderm (*EOMES*), mesoderm (*MESP1*), cardiogenic mesoderm (*TNNI1*), endoderm (*SOX17*), and mesoderm-derived endothelial cells (*PECAM1*). (**I**) Cluster distribution of cells from all experimental perturbations (compare UMAP in **Fig. 5A**). Note that [*SOX2↓ NANOG↓*] powerfully pushes iPSCs out of the pluripotent state and toward mesoderm and endoderm using these experimental conditions. Note, too, that differentiation is less efficient with [*TBX6*↑ *SP5*↑] than [*TBX6*↑]. We later found this effect likely to be caused by dilution of available dCas12a-miniVPR protein (see **Fig. S6J**), and could be corrected by boosting Cas gene expression levels (see **Fig. S12A-B**). (**J**) The reason that addition of [*SP5*↑] leads to repression of endothelial cells is likely that *SP5* is expressed by early mesoderm and endoderm but not endothelial cells (red dashed line). (**K**) PreciCE algorithm output for endoderm to mesoderm, as used to actively block endoderm formation. (**L**) Parameter plot for this conversion. (**M-N**) Expression of *HNF1B* and *MESP1* are restricted to endoderm and mesoderm, respectively, allowing us to use *HNF1B* and *MESP1* as RT-qPCR markers to analyze differentiation. All error bars represent standard deviation.

**Figure S11.**
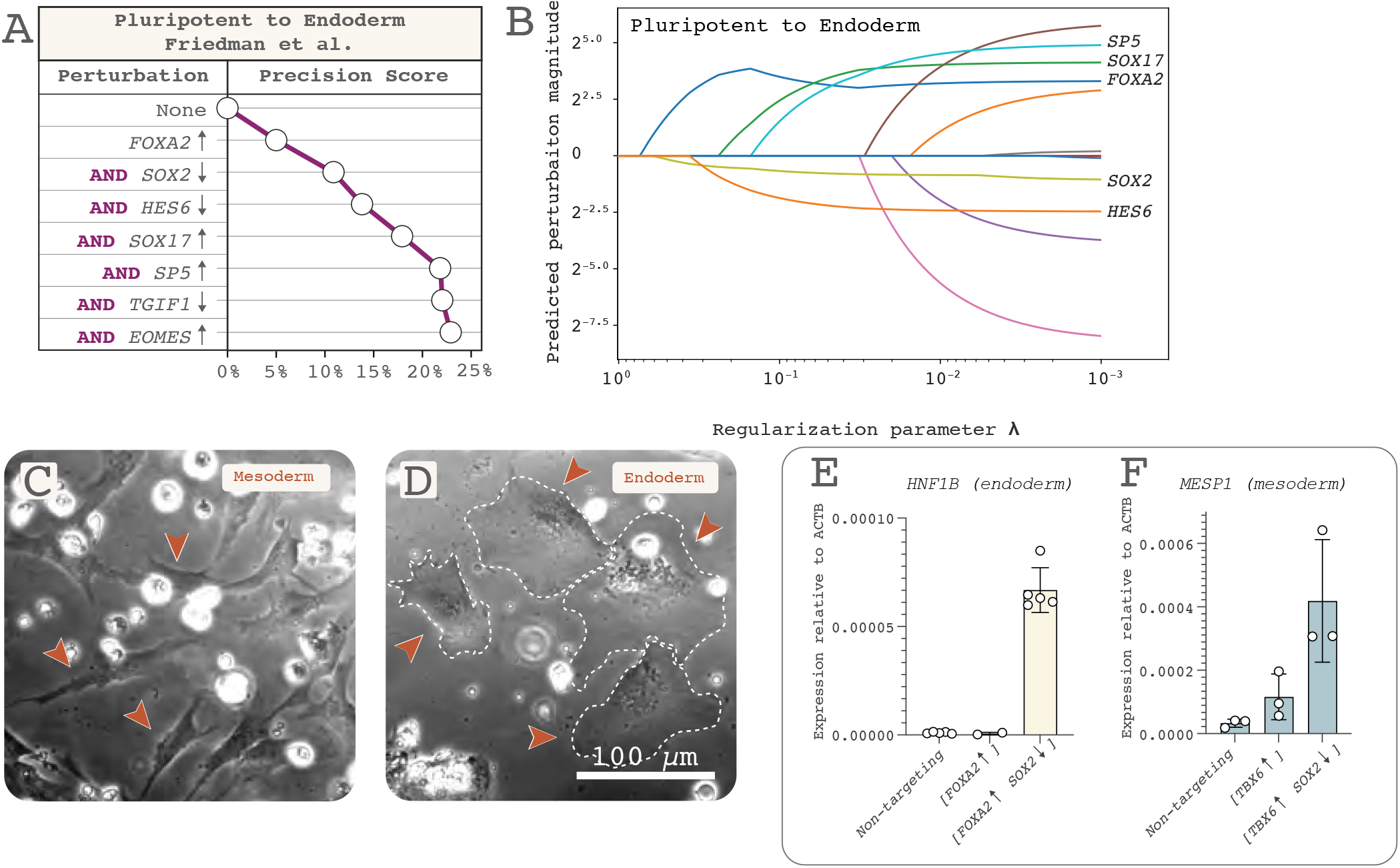
Cell fate logic using PreciCE for endoderm differentiation. (**A-B)** PreciCE algorithm output for the conversion of pluripotent stem cells to endoderm (**A**) and associated parameter plot (**B**). (**C-D**) Mesoderm (**C**, through [*TBX6*↑ *SP5*↑ *SOX2↓ FOXA2↓ SOX17↓*]) and endoderm cells (**D,** through [*FOXA2*↑ *SOX2↓*]) differ dramatically in their morphology, the latter displaying a large, flattened appearance (arrowheads show example cells; dashed lines show cell outlines; white round dots are floating dead cells). (**E**) [*FOXA2*↑] can only execute iPSC-to-endoderm conversion if it is combined with repression of the starting state ([*SOX2↓*]). (**F**) This differs from iPSC-to-mesoderm conversion, where [*TBX6*↑] generates mesoderm (albeit inefficiently) even in the absence of [*SOX2↓*]. Note that panels **E** and **Fig. 5I** came from one single experiment so the data for the non-targeting condition are the same for panels **E** and **Fig. 5I**. All error bars represent standard deviation.

**Figure S12.**
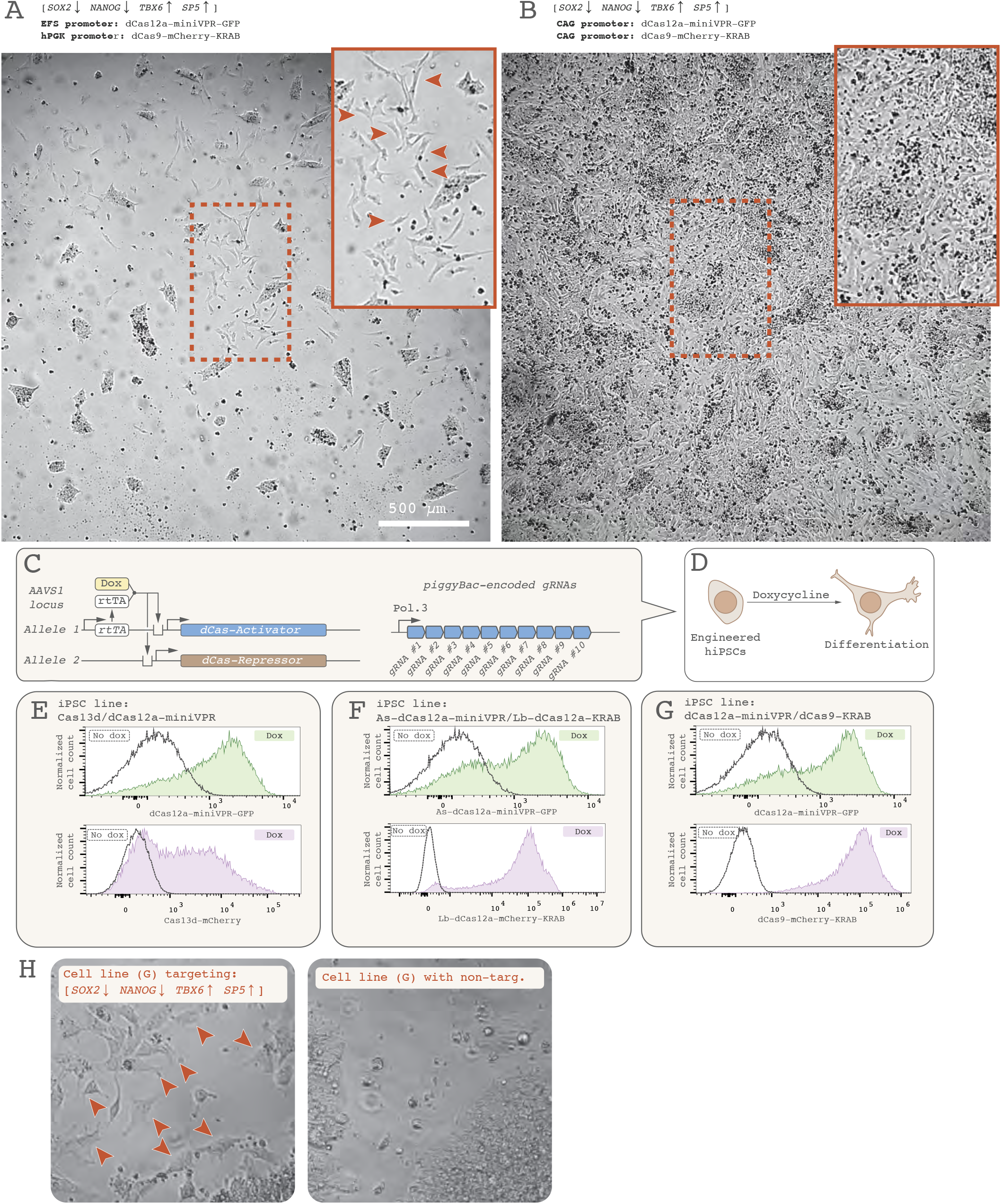
Mesoderm differentiation using an engineered iPSC line, and dependence on Cas gene expression level. (**A**) Cell fate conversion efficacy is dramatically improved by replacing the weak EFS and hPGK promoters driving dCas12a and dCas9, respectively with a strong CAG promoter (**B**). Compare the scattered mesoderm cells in (**A**) with the densely packed carpet of mesoderm cells in (**B**; rectangles show inset from dashed line). This shows that increased Cas gene expression strongly determines efficacy of gene modulation and cell differentiation. **(C)** Engineered iPSC lines carry targeted genomic insertions of Cas genes into the *AAVS1* safe-harbor locus (one construct per *AAVS1* allele). Constitutive expression of a reverse tetracycline transactivator (rtTA) enables doxycycline-inducible expression of a dCas12a-miniVPR-GFP activator and a dCas9-mCherry-KRAB repressor. gRNA constructs are inserted in this cell line through piggyBac-mediated integration. (**D**) With these cell lines, cell differentiation is triggered through the addition of doxycycline to the culture medium. (**E-G**) iPSC lines carrying the Cas13d/dCas12a system (**E**), As-dCas12a/Lb-dCas12a system (**F**), and dCas9/dCas12a system (**G**) activate Cas gene expression when treated with doxycycline (dark trace represent same cell lines not treated with doxycycline). (**H**) In the dCas9/dCas12a cell line (**G**), doxycycline administration triggers mesoderm differentiation, as shown by the morphological changes (arrowheads) compared to the cells expressing non-targeting gRNAs.

